# Genetic mapping reveals new loci and alleles for flowering time and plant height using the double round-robin population of barley

**DOI:** 10.1101/2023.01.12.523733

**Authors:** Francesco Cosenza, Asis Shrestha, Delphine Van Inghelandt, Federico A. Casale, Po-Ya Wu, Marius Weisweiler, Jinquan Li, Franziska Wespel, Benjamin Stich

## Abstract

Flowering time and plant height are two critical determinants of yield potential in barley (*Hordeum vulgare*). Although their role as key traits, a comprehensive understanding of the genetic complexity of flowering time and plant height regulation in barley is still lacking. Through a double round-robin population originated from the crossings of 23 diverse parental inbred lines, we aimed to determine the variance components in the regulation of flowering time and plant height in barley as well as identify new genetic variants by single and multi-population quantitative trait loci (QTL) analyses and allele mining. Despite similar genotypic variance, we observed higher environmental variance components for plant height than flowering time. Furthermore, we detected one new QTL for flowering time and two new QTL for plant height. Finally, we identified a new functional allelic variant of the main regulatory gene *Ppd-H1*. Our results show that the genetic architecture of flowering time and plant height might be more complex than reported earlier and that a number of undetected, small effect or low frequency, genetic variants underlie the control of these two traits.

## 1 INTRODUCTION

The increase in world population, the reduction of available arable land, and climate change represent some of the greatest challenges that humanity is and will further have to face in the near future (Vyas *et al*., 2022). One of the answers to these challenges is to reduce the influence of biotic and abiotic stress factors and by that increase crop productivity (Khush, 2013). Of particular importance are yield increases of cereals (Araus *et al*., 2008), which are essential for human nutrition as they alone contribute about 44.5% of the calory uptake of the world population (FAO, 2019). In addition, they are important for animal feeding and beverage production (FAO, 2020).

Flowering time is one of the critical determinants of yield potential in cereals (Hill and Li, 2016). This is because, at this phenological stage, the plant transits from the vegetative to the reproductive phase, and grain filling starts (Cockram *et al*., 2007). In turn, the efficiency of grain filling is maximized if it coincides with optimal environmental conditions (Wiegmann *et al*., 2019). Therefore, plants and farmers have adapted several strategies to synchronize the phenological stages to environmental conditions (Anderson and Song, 2020).

Barley (*Hordeum vulgare* L.) is ranked fourth among the most cultivated cereals worldwide (FAO, 2020). This species is characterized by great environmental plasticity that allows it to be cultivated at different latitudes, with extremely dissimilar conditions of temperature and photoperiod (Dawson *et al*., 2015). It has been demonstrated that the adaptive success of barley is also due to the selection of favorable allelic variants at the main genes determining the transition from the vegetative to the reproductive phase (Comadran *et al*., 2012; Göransson *et al*., 2019; Turner *et al*., 2005). Three types of genes have been identified to be responsible for the modulation of flowering time in barley: genes that act under the influence of photoperiod, genes that act under the influence of temperature, and genes, called as *earliness per se*, that act independently of environmental variables (Fernández-Calleja *et al*., 2021).

The main genes whose expression is influenced by the photoperiod are *Ppd-H1* (Turner *et al*., 2005) and *Ppd-H2* (Kikuchi and Handa, 2009). *Ppd-H1*, which is located on chromosome 2H, has been described as the major determinant of the response to long day conditions in barley, acting jointly with *HvCO1* and *HvCO2* (Campoli *et al*., 2012). At the same time, *Ppd-H1* indirectly influences the response to vernalization by promoting the expression of *Vrn-H3* (Mulki and von Korff, 2016). *Ppd-H2* is the second main driver of the photoperiod response in barley, but unlike *Ppd-H1* it acts in short day conditions. The non-functional allelic variant of *Ppd-H2* allowed the expansion of the cultivation area of barley at higher latitudes (Casao *et al*., 2011).

The major determinants of the response to temperature are genes involved in the vernalization process. *Vrn-H1*, which is located on chromosome 5H, promotes flowering after the plant has satisfied its vernalization requirement (Yan *et al*., 2003). Furthermore, *Vrn-H1* inhibits the expression of *Vrn-H2*, which is located on chromosome 4H. *Vrn-H2* delays flowering, allowing the plant to fulfill its cold needs (Deng *et al*., 2015; Yan *et al*., 2004). The interaction between *Vrn-H1* and *Vrn-H2* is therefore one of the main mechanisms that allow the control of flowering time in winter or facultative barley varieties (Yan *et al*., 2004). The third gene responsible for the temperature response in barley is *Vrn-H3* on chromosome 7H (Yan *et al*., 2006). *Vrn-H3*, when not repressed by *Vrn-H2*, promotes flowering by allowing the transition from the vegetative to the reproductive phase in long day conditions (Hemming *et al*., 2008).

Within the group of *earliness per se* genes, the major determinant is *HvCEN* which is located on chromosome 2H. Because its expression is not directly influenced by environmental variables, the allelic variants of *HvCEN* allowed the expansion and adaptation of barley to new areas through the regulation of flowering time (Comadran *et al*., 2012). In addition, three other genes have been described as circadian clock-related *earliness per se* genes which, although not directly influencing flowering, alter the expression of *Ppd-H1*: *HvELF3* (Faure *et al*., 2012), on chromosome 1H, *HvLUX1* (Campoli *et al*., 2013), on chromosome 3H, and *HvPHYC* (Nishida *et al*., 2013), on chromosome 5H. Furthermore, mutations in *HvELF3* can also affect the expression of *HvGI* (Dunford *et al*., 2005), causing earlier flowering (Zakhrabekova *et al*., 2012). Finally, other genes initially reported to be responsible for controlling other quantitative traits have also been described to have an influence on flowering time or flower development: *HvAP2* (Shoesmith *et al*., 2021), on chromosome 2H, and *Hv20ox2* (*sdw1/denso*) (Bezant *et al*., 1996; Jia *et al*., 2009), on chromosome 3H. Another key trait responsible for determining production performance in cereal species is plant height (Mikołajczak *et al*., 2017). An adequate plant height allows to obtain a lower exposure to lodging and a higher harvest index but on the other side, it is essential to keep the spikes far from the soil to reduce the risk of yield losses caused by infectious diseases (Vidal *et al*., 2018). Plant height and flowering time are two interrelated characters. This is because flowering is possible when the meristem has switched from the vegetative to the reproductive phase. For this reason, many of the genes controlling flowering time, such as *Ppd-H1* (Turner *et al*., 2005), *Vrn-H1* (Wiegmann *et al*., 2019), *Vrn-H2* (Rollins *et al*., 2013), *Vrn-H3* (Arifuzzaman *et al*., 2016), *Hv20ox2* (Jia *et al*., 2009), *HvCEN* (Bi *et al*., 2019), and *HvAP2* (Patil *et al*., 2019), have a pleiotropic effect on plant height. In addition to these genes, other genes involved in the biosynthesis of brassinosteroids, such as *HvBRD* on chromosome 2H, *HvBRI1* (*uzu*) on chromosome 3H, *HvDWF4*, on chromosome 4H, *HvCPD* and *HvDEP1* on chromosome 5H, and *HvDIM* on chromosome 7H (Dockter *et al*., 2014; Wendt *et al*., 2016), have been described to be involved in plant height regulation of barley. Some of the above mentioned genes, such as *HvAP2* and the genes regulating brassinosteroids biosynthesis, have been identified based on mutant approaches (Dockter *et al*., 2014; Shoesmith *et al*., 2021). When instead natural variation was exploited, bi-parental (Arifuzzaman *et al*., 2014; Von Korff *et al*., 2006; Rollins *et al*., 2013; Schmalenbach *et al*., 2009) or nested association mapping populations (Maurer *et al*., 2015; Nice *et al*., 2017) were used. When multi-parental populations were examined instead, the experiments comprised a restricted number of inbred lines (Cuesta-Marcos *et al*., 2008), and/or the selected parental inbreds were from a restricted geographical range (Afsharyan *et al*., 2020; Cuesta-Marcos *et al*., 2008). All these factors reduce the likelihood of identifying genes and allelic variants with low population frequency (Yu *et al*., 2006). Therefore, the utilization of segregating populations derived from genetic resources with high genotypic and phenotypic diversity could allow the identification of further genes that are mechanistically involved in flowering time and plant height regulation. This has the potential to facilitate and speed up breeding and provide new targets for genetic modification through, for example, CRISPR platforms. In turn, this could help to extend the cultivation area of barley by allowing its adaptation to new environmental conditions. Furthermore, the knowledge gained in barley has a high potential to be transferred to other cereal species that are genetically close but have a polyploid chromosomal structure, such as tetraploid (*Triticum turgidum* var. *durum*) and hexaploid (*Triticum aestivum*) wheat (Langridge, 2018).

In this study, a multi-parent population was used to explore the genetic landscape of flowering time and plant height in barley with the objectives of: (i) determining the genetic variance components in the regulation of flowering time and plant height, (ii) obtaining a comprehensive understanding of the genetic complexity of flowering time and plant height in barley by single and multi-population QTL analyses, (iii) identifying candidate genes for the detected QTL regulating flowering time and plant height, and detecting new allelic variants of genes responsible for the control of these two traits.

## 2 MATERIALS AND METHODS

### 2.1 Plant material and genotypic evaluation

The plant material used in this study consisted of a population which is designated in the following as *Hordeum vulgare* Double Round-Robin (HvDRR). The population originated from the crossings of 23 parental inbred lines, including eleven cultivars and twelve landraces (Shrestha *et al*., 2022), in a double round-robin scheme (Stich, 2009) (Supplementary Table 1). The parental inbred lines have been chosen from a diversity panel of 224 spring barley accessions selected from the Barley Core Collection (BCC) (Pasam *et al*., 2012) to maximize the combined genotypic and phenotypic richness index (Weisweiler *et al*., 2019).

Starting from the 45 F1s, a single seed descent strategy has been applied to develop between 35 and 146 recombinant inbred lines (RIL) for each of the 45 sub-populations (Casale *et al*., 2022). The RILs were genotyped at the F4 generation using a 50K SNP genotyping array (Bayer *et al*., 2017).

### 2.2 Phenotyping

Flowering time (FT) evaluation was carried out in seven environments in Germany: Cologne from 2017 to 2019, Mechernich from 2018 to 2019, and Quedlinburg from 2018 to 2019. Plant height (PH) was evaluated in the same environments except for Quedlinburg, totaling five environments. At the Cologne and Mechernich environments, 33 seeds were sown in single rows of 1.6 meters length. In Quedlinburg, double rows of the same length were sowed. The inter-row distance was 20 cm. Fertilization and plant protection followed local practices.

In each environment, an augmented design was used. RILs of the HvDRR population and the inbreds of the diversity panel were planted with one replicate and only the parental inbreds of the HvDRR population were replicated 15-20 times per environment. FT was recorded as days after sowing when 50% of the plants within the (double) row were flowering. PH was measured on average across all available plants within a row as height in cm from the collar to the peak of the plant when the spike was fully developed.

### 2.3 Statistical analyses

The collected phenotypic data were subject to statistical analysis using the following linear mixed model:

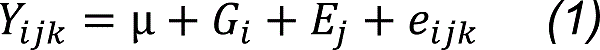

where *Y_ijk_* indicated the observed phenotypic value for the *i^th^* genotype in the *j^th^* environment within the *k^th^* replication, *μ* the general mean of the trait, *G_i_* the effect of the *i^th^* genotype, *E_j_* the effect of the *j^th^* environment, and *e_ijk_* the random error. For the calculation of adjusted entry means, the genotypic effect was considered fixed, while the environmental effect was considered random.

The broad sense heritability (*h^2^*) was calculated as:

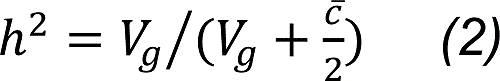

where *V_g_* represented the genotypic variance and ĉ the mean of the standard errors of the contrasts among all pairs of genotypes (Piepho and Möhring, 2007). For the calculation of the genotypic variance (*V_g_*), model *(1)* was used, but all effects were considered as random. In addition, we calculated *h^2^*, when applying for each environment a correction based on the augmented design considering different grid sizes, and then estimating *V_g_* and ĉ across the environments.

In order to quantify the interaction between genotype and environment, we used a second linear mixed model:

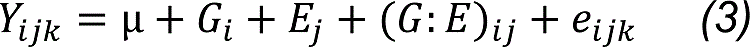

where (*G*:*E*)_ij_ represented the interaction between the *i^th^* genotype in the *j^th^* environment, which was fitted to the data of the parental inbreds.

#### 2.3.1 QTL analyses

Two different QTL analyses were performed in this study on the HvDRR population: multi-parent population (MPP) and single population (SP) analyses.

The estimation of genetic maps necessary for the SP analysis, as well as that of the consensus map used in the MPP, have been described by Casale *et al*. (2022).

For each sub-population and each trait, an SP QTL analysis was performed, based on the adjusted entry means for each RIL calculated with model *(1)*, using the following scheme: first, standard interval mapping using the Haley-Knott regression algorithm (Knott and Haley, 1992) was applied, followed by forward selection in order to determine the number of QTL to include in the model. Then a forward and backward selection algorithm was applied to perform multiple QTL mapping. Model selection was based on the highest penalized LOD score with penalties determined through 4000 permutations. A two-dimensional genome-wide scan was performed to detect epistatic interactions between all pairs of loci in the genome. The SP analyses were carried out with the R package “qtl” (Broman *et al*., 2003). The MPP analyses were performed by jointly analyzing all sub-populations using an ancestral model that took into account the degree of relatedness among the parental inbreds (Garin *et al*., 2017). The degree of relatedness was calculated by clustering the haplotypes. The haplotype window size was chosen as the consensus genetic map distance for which the linkage disequilibrium (LD), measured as *r²*, was 0.2 (Giraud *et al*., 2014) (Supplementary Table 2). The MPP analysis was performed using the R package “mppR” (Garin *et al*., 2015).

Confidence intervals for the QTL detected via SP and MPP were calculated using a 1.5 LOD drop method (Manichaikul *et al*., 2006).

#### 2.3.2 Genomic prediction

Genomic predictions of FT and PH in the HvDRR population were performed by genomic best linear unbiased prediction (GBLUP) using the following model (VanRaden, 2008):

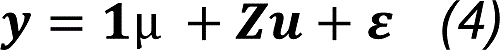

where ***y*** was the vector of the adjusted entry means of the considered trait (FT or PH), **1** was a unit vector, *μ* the general mean, ***Z*** the design matrix that assigned the random effects to the genotypes, and ***u*** the vector of genotypic effects that were assumed to be normally distributed with *N*(0, *K*σ_u_^2^), in which ***K*** denoted the realized kinship matrix between inbreds and σ_u_^2^ the genetic variance of the GBLUP model. In addition, ***ε*** was the vector of residuals following a normal distribution *N*(0, Iσ_e_^2^).

The prediction ability of the GBLUP model was evaluated by Pearson’s correlation coefficient (*r*) between observed and predicted phenotypes. To assess the model performance, five-fold cross-validation (CV) with 20 replications was performed. In that case, the prediction ability was defined as the median of the prediction abilities across the 20 runs of each 5 fold-CV.

#### 2.3.3 Candidate gene analysis and allele mining

The candidate gene analysis was performed for those QTL from the SP analysis that did not carry inside their confidence intervals previously reported genes controlling the corresponding trait, explained ≥ 15% of the phenotypic variance, and had a confidence interval ≤ 30 cM. For the QTL that fulfilled these criteria, all the genes within the confidence interval were extracted using the Morex v3 reference sequence (Mascher *et al*., 2021). Next, variant calling data of single nucleotide polymorphisms (SNPs), causing tolerated and deleterious mutations, insertion and deletions (INDELs), and predicted structural variants (SV), obtained as described by Weisweiler *et al*. (2022), were used to identify genes that were polymorphic between the two parental inbreds of the sub-population in which the QTL was detected. For each gene, we took into account all the polymorphisms inside the coding, non-coding, and, for SV, potential regulatory regions of the gene within 5 kb up and downstream of the gene.

Subsequently, we performed a weighted gene co-expression network analysis (WGCNA) to identify modules of co-expressed genes that were associated with the phenotypic variability of the traits. The mRNA sequencing experiment of leaf samples of 21 parental inbred lines, described by Weisweiler *et al*. (2019), was the basis for this analysis. The selected soft thresholding power was two, based on the scale-free topology criterion (Zhang and Horvath, 2005). We predicted the gene networks for the three modules with the highest and the three lowest correlations for both traits. In order to have a comprehensive understanding of the networks we selected genes with a gene-module membership p-value < 0.01 and, within them, the top 30% of gene-gene interactions based on the interactions weight. Because of the high number of gene-gene interactions in the module “turquoise”, we selected the top 5% of interactions with the highest weight. For the “lightyellow” and “tan” modules we did not filter the interactions based on weight. Furthermore, for the “black” module we selected the genes with a gene-module membership p-value < 0.05.

In the next step, the results of the WGCNA and SP QTL analyses were combined: we further filtered the polymorphic genes within the confidence intervals based on their membership to a module (Wei *et al*., 2022). The genes within the three modules with the highest and the three with the lowest correlation with the trait under consideration were evaluated for their functional annotation. We selected as candidate genes those with an annotation similar to that of genes previously reported to control the trait under consideration in barley and those for whom functional annotation has been described to be involved in plant vegetative or reproductive development. All the analyses for the calculation of the weighted gene co-expression networks were performed with the R package “WGCNA” (Langfelder and Horvath, 2008).

Allele mining was performed for the known genes with pleiotropic effect on FT and PH located within SP QTL confidence intervals. For each gene, polymorphisms between the parental inbreds of the respective segregating sub-populations were extracted from the whole genome sequencing data (Weisweiler *et al*., 2022). To confirm the accuracy of the whole genome sequencing data, we performed Sanger sequencing of the 23 parental inbreds for *Ppd-H1*. To predict the effect of polymorphisms on the phenotype, we used the SIFT algorithm (Vaser *et al*., 2016). In addition, we performed PCR, as described in Karsai *et al*. (2005), to check the presence/absence of the three *Vrn-H2* genes.

#### 2.3.4 Fine mapping of QTL by association genetics

We used association genetics in the diversity panel of Pasam *et al*. (2012) to fine map the QTL that did not carry within their confidence intervals genes reported to control the corresponding trait, explained ≥ 15%, and had a confidence interval ≤ 30 cM. We used the phenotypic data of the 224 inbreds collected in our field trials and the genotypic information available from Milner *et al*. (2019). To construct the kinship matrix among the 224 inbreds, we used all the SNPs in the SNP matrix. Association analysis was performed using only polymorphisms from QTL fulfilling the above mentioned criteria. For association analysis, we used a mixed model approach, implemented for the variance component (Kang *et al*., 2010), with the R package “statgenGWAS” (van Rossum *et al*., 2022).

## 3 RESULTS

### 3.1 Phenotypic variation and covariation

FT and PH were evaluated for each RIL across seven and five environments, respectively. For both traits, the environmental variance (*V_e_*) was about two to three times higher than the genotypic variance (Table 1). Furthermore, for FT, the G:E variance was about half the genotypic variance, while for PH the G:E variance was about 87% of the genotypic variance. The values of broad-sense heritability on an entry means basis were high to very high, ranging from 0.76 for PH to 0.86 for FT (Table 1).

**Table 1:** Variance components of the multi-environment linear mixed model and heritability values for flowering time and plant height. G represents the genetic variance, E the environmental variance, G:E the variance explained by the interaction between G and E, and e the residual error.

| Trait | Groups | Variance | $h^2$ |
| --- | --- | --- | --- |
| Flowering time | G | 41.33 | 0.86 |
|  | E | 77.12 |  |
|  | G:E | 22.31 |  |
|  | e | 17.02 |  |
| Plant height | G | 41.46 | 0.76 |
|  | E | 128.55 |  |
|  | G:E | 36.32 |  |
|  | e | 56.04 |  |

To take into account possible intra-environmental variation, the phenotypic values were adjusted using moving grids of three different sizes, exploiting the possibilities of an augmented design. For all three examined grid sizes, we observed that the resulting heritability values across all environments were reduced compared to the analysis without adjustment. Therefore, we decided to discuss in the following only results from analyses where intra-environmental variation was not corrected for.

Across all environments, the sub-population that was found to be the earliest to flower was HvDRR35, where RILs flowered on average 58 days after sowing. In contrast, the latest sub-population to flower was HvDRR46 for which, on average, RILs reached flowering 79 days after sowing (Figure 1; Supplementary Table 3). HvDRR46 was also the sub-population with the smallest plants, with an average height of 48 cm. In contrast, HvDRR12 was, with an average of 87 cm, the sub-population with the tallest plants (Figure 1; Supplementary Table 3). HvDRR09 was the sub-population with the lowest coefficient of variation (CoV) value for FT (2.73 days), while the highest CoV was observed for HvDRR43 (23.31 days) (Supplementary Table 3). Regarding PH, the sub-population with the smallest variability was HvDRR15 (CoV = 3.83 cm), while the highest CoV, 30.53 cm, was observed for HvDRR46 (Supplementary Table 3). The CoV was, for the diversity panel across the same environments, 7.35 days for FT and 14.01 cm for PH (Supplementary Table 3).

**Figure 1:**
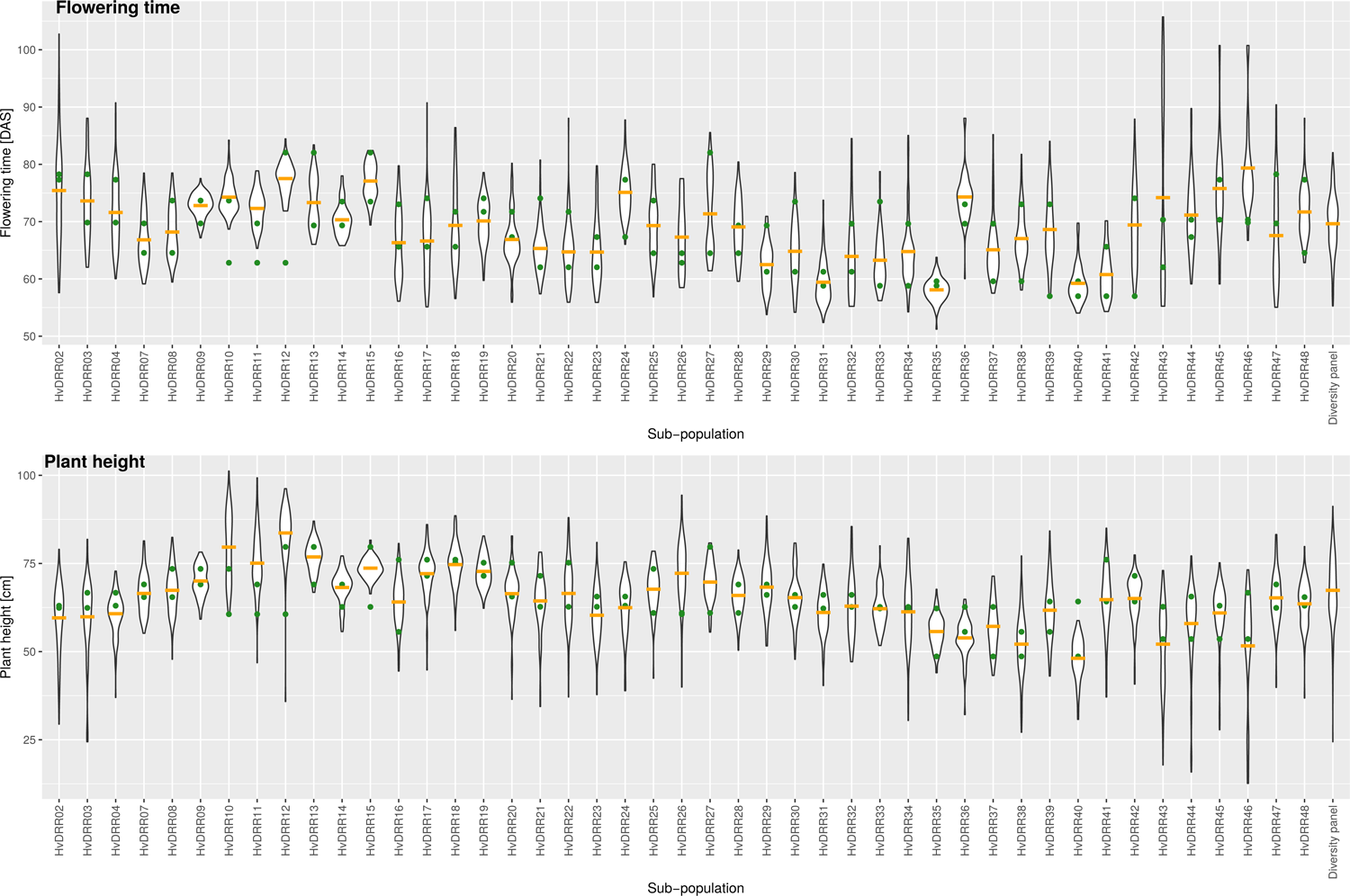
Violin plots for adjusted entry means for flowering time and plant height of each HvDRR sub-population and for the 224 inbreds of the diversity panel. The flowering time is presented as days after sowing (DAS) and the plant height values are reported in cm. The green dots represent the adjusted entry means of the parental inbreds of the sub-population. The orange lines represent the average of the adjusted entry means of the recombinant inbred lines of the respective sub-population.

The differences between the mean of the parental inbreds and the mean of the sub-populations were also examined as these are an indicator for the presence of epistasis. For flowering time, the differences between the means of the parental inbred lines and the respective sub-populations were statistically significant (p < 0.05) in 22 cases. Among them, the highest differences were observed for sub-populations HvDRR43, with 7.2 days, and HvDRR46, with 10.5 days. For plant height the differences of the means of the sub-populations and the parental inbreds were significant (p < 0.05) in 14 cases. The strongest difference between the parental inbreds and the progeny mean was observed for sub-populations HvDRR10, with 9.62 cm, HvDRR12, with 9.88 cm, and HvDRR11, with 10.60 cm. All these sub-populations had Sanalta as common parental inbred (Figure 1).

Across all sub-populations, the correlation coefficient of FT and PH was −0.012 (Supplementary Figure 1). However, considerable differences were observed for the single sub-populations (Figure 2). HvDRR28 was the sub-population for which the highest correlation coefficient has been observed (0.44), while the sub-population where the two traits were most negatively correlated was HvDRR43 (−0.77) (Supplementary Figure 2).

**Figure 2:**
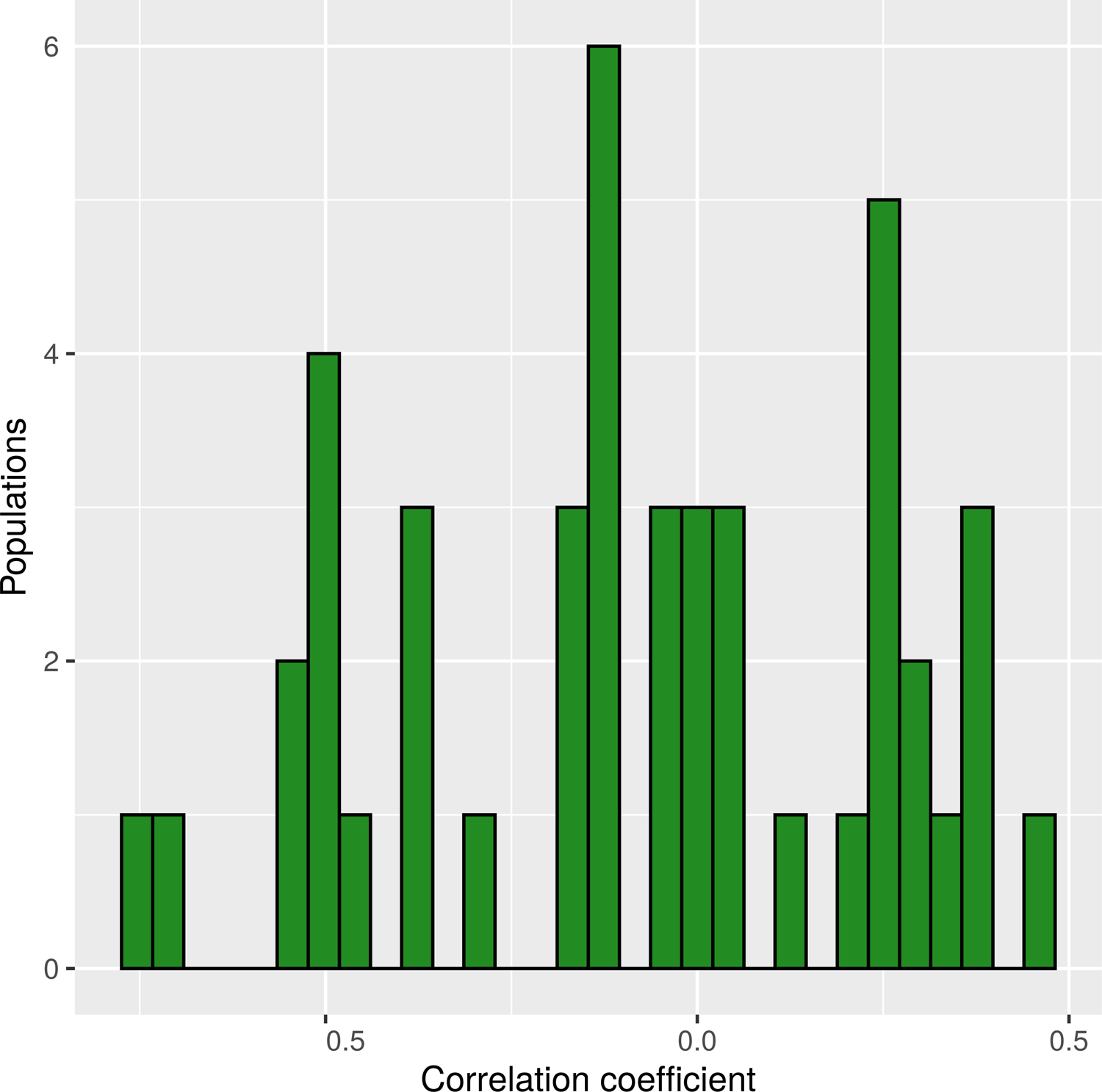
Distribution of correlation coefficients between flowering time and plant height calculated for the HvDRR sub-populations. On the x axis the correlation coefficients are represented and on the y axis the number of sub-populations.

### 3.2 Multi-parent population analysis

The multi-parent population analysis identified each 21 QTL for FT and PH, distributed across all seven chromosomes (Figure 3). The analysis was performed using the genetic haplotype window sizes estimated from the extent of linkage disequilibrium (Supplementary Table 2). The percentage of phenotypic variance explained by all the QTL detected in the MPP analysis was 39.1% and 24.9% for FT and PH, respectively. For FT, the confidence interval of 17 QTL overlapped with the interval of at least one QTL identified in the SP analysis (Supplementary Tables 4-5). Out of 21 QTL identified for PH, 16 overlapped with one or more QTL detected in SP analysis (Supplementary Tables 4-6). Among the QTL detected for both traits, the intervals of two pairs of QTL overlapped: *FT-MP-Q3* with *PH-MP-Q3* and *FT-MP-Q19* with *PH-MP-Q20*.

**Figure 3:**
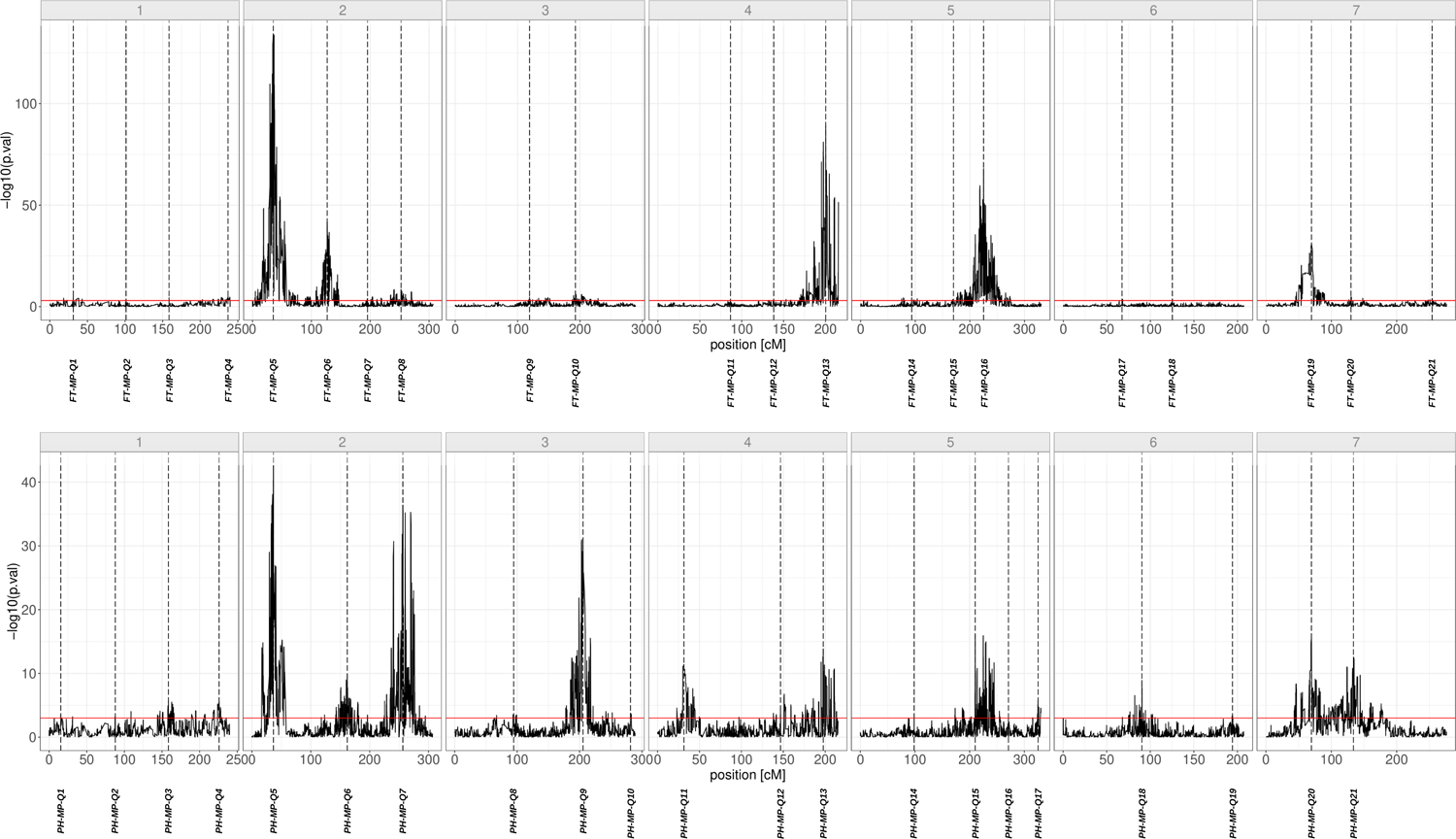
Negative decadic logarithm of the p-value of the multi-parent population analysis for flowering time (top) and plant height (bottom) using an ancestral model. On the x axis, the position on the consensus genetic map is reported. Each dashed line indicates the peak position of the corresponding QTL.

The additive effect of the 23 parental inbreds for the 21 QTL for FT ranged from −23.42 days, observed for Ancap2 at *FT-MP-Q5,* to 5.14 days, for Kombyne at *FT-MP-Q13* (Figure 4). However, in about 92% of cases, the additive effect for FT was between −1 and 1 day (Supplementary Figure 3). For PH, the effect ranged from −3.88 cm for Kombyne at *PH-MP-Q15*, to a maximum of 1.99 cm at *PH-MP-Q5*, for seven parental inbreds (Figure 4). Also in this case more than 90% of the additive effects had a value between −1 and 1 cm (Supplementary Figure 3). The crossings design underlying our population allows to estimate the number of alleles at each QTL. The QTL with the highest number of significantly different allele effects and thereby with presumably alleles were, for FT, *FT-MP-Q20*, with 9 alleles, and, for PH, *PH-MP-Q20*, with 8 detected alleles.

**Figure 4:**
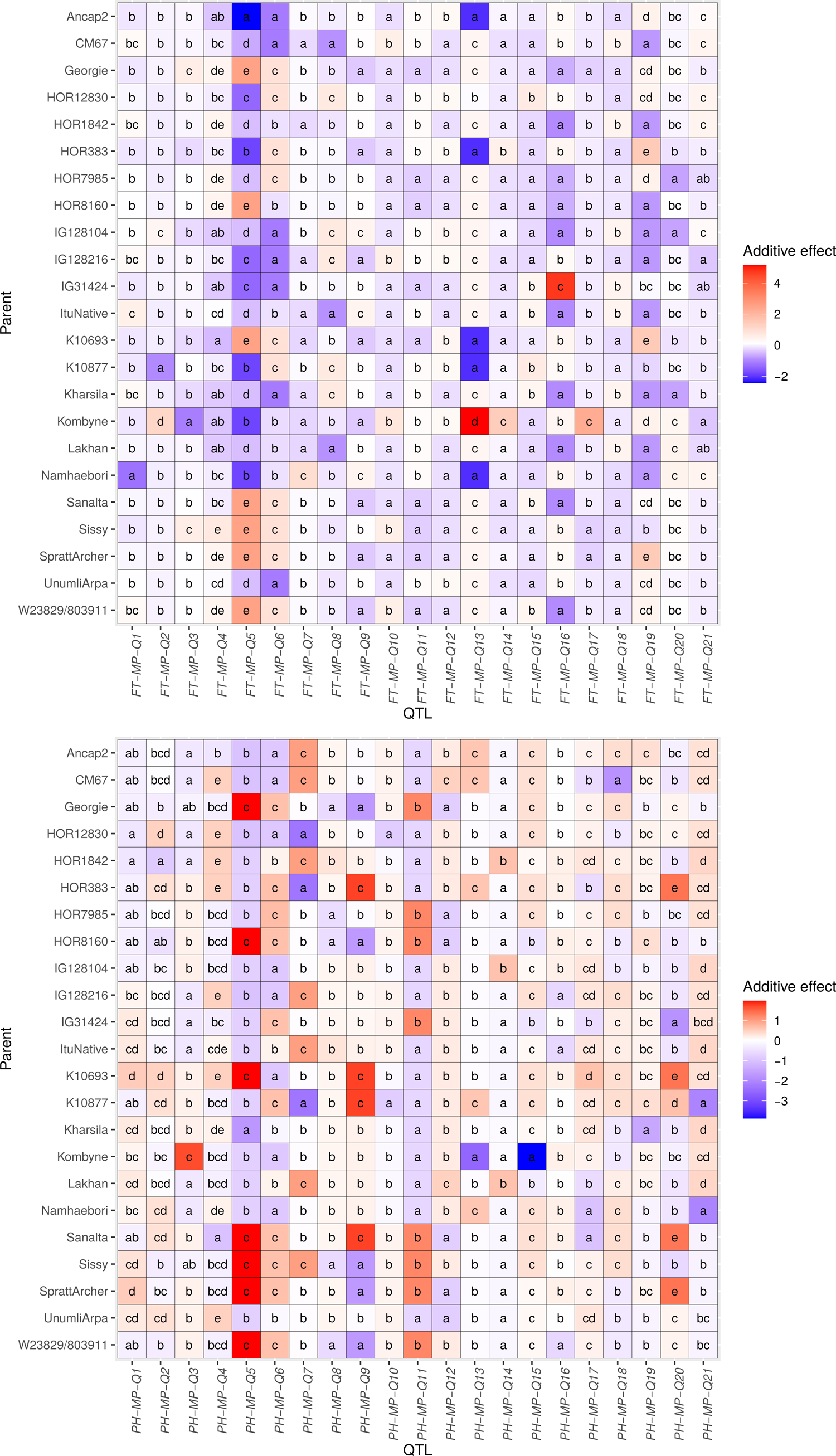
Heat map of the effects of the parental inbreds at the QTL detected through multi-parent population analysis for flowering time (top, in days after sowing) and for plant <u>height</u> (bottom, in cm). Indexed letters indicate the significance of the difference (p < 0.05) between the effects of the same QTL.

### 3.3 Genomic prediction ability

The prediction abilities of the GBLUP model across the HvDRR population were high with values of 0.89 and 0.87 for FT and PH respectively (Supplementary Table 7). To compare the prediction performance of the GBLUP model with those of the detected QTL, we used the squared prediction abilities. The coefficient of determinations (*r²*) obtained by genomic prediction without CV was 0.79 for FT and 0.76 for PH. The cross-validated prediction abilities were 0.77 for both FT and PH.

### 3.4 Single population QTL analysis

Through single population analysis, 89 QTL were identified for FT and 80 for PH (Figures 5-6; Supplementary Tables 5-6). The percentages of explained variance by the individual QTL detected for FT sized from 1.02%, for *qHvDRR47-FT-7.1*, to 77.75%, for *qHvDRR27-FT-2.1* (Supplementary Table 5), while for PH the values ranged from 2.52%, for *qHvDRR11-PH-2.2*, to 63.62%, for *qHvDRR10-PH-3.1* (Supplementary Table 6). HvDRR22 was the sub-population with the highest values of explained variance for both FT (70.24%) and PH (78.54%), while the lowest percentages of variance explained by the detected QTL were observed for FT in HvDRR35 (33.49%) and for PH in HvDRR31 (23.65%) (Supplementary Tables 5-6).

**Figure 5:**
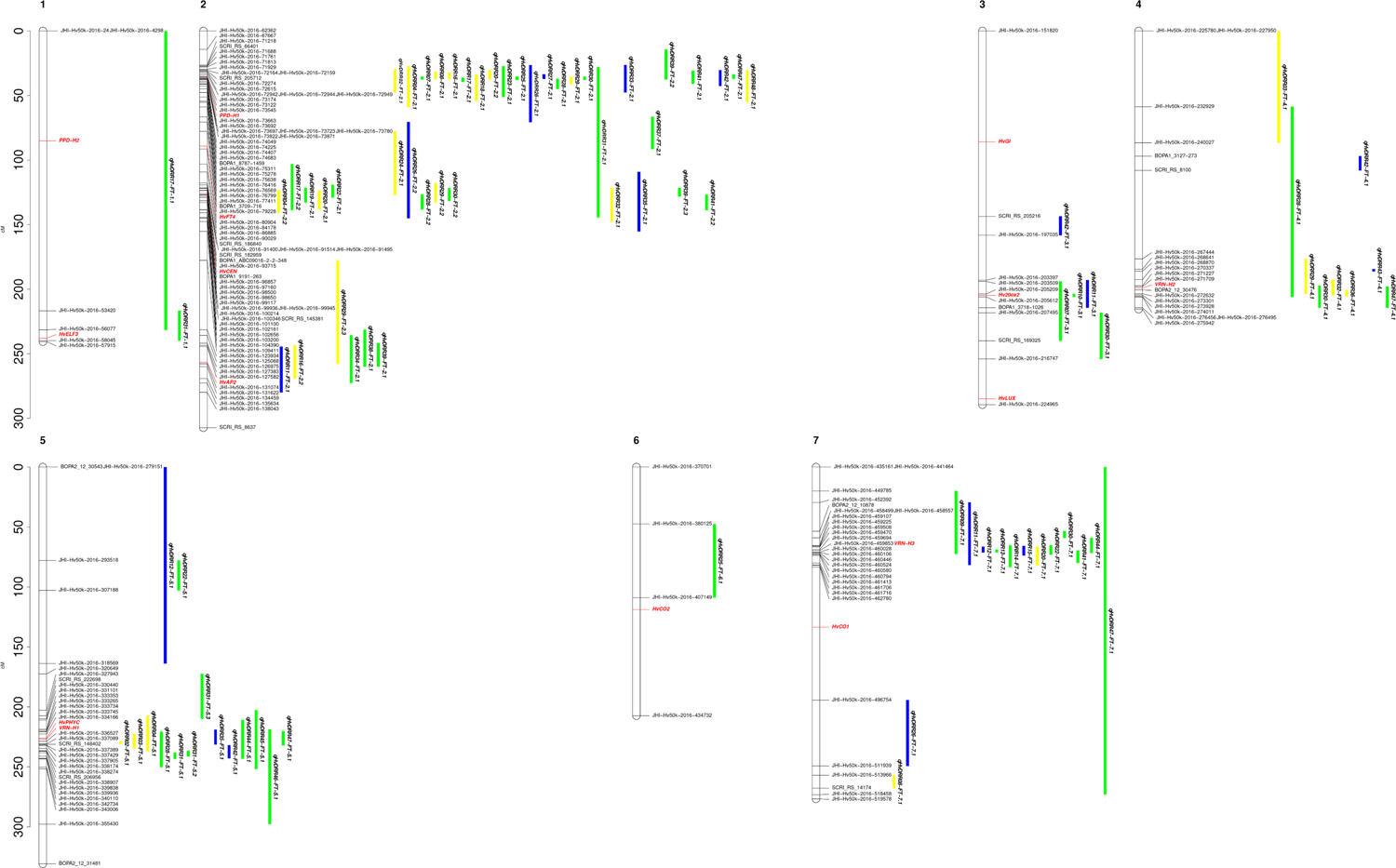
Genetic position of the QTL detected in single population analyses for flowering time projected to the consensus map. The position of the QTL confidence intervals is represented as a vertical bar parallel to the right of the chromosome. The color of the bar indicates if the sub-population was obtained by crossing two landraces (yellow), two cultivars (blue), or a landrace and a cultivar (green). The genetic positions of the known genes regulating flowering time in barley are shown in red. The positions of the markers that flank each QTL are also reported.

**Figure 6:**
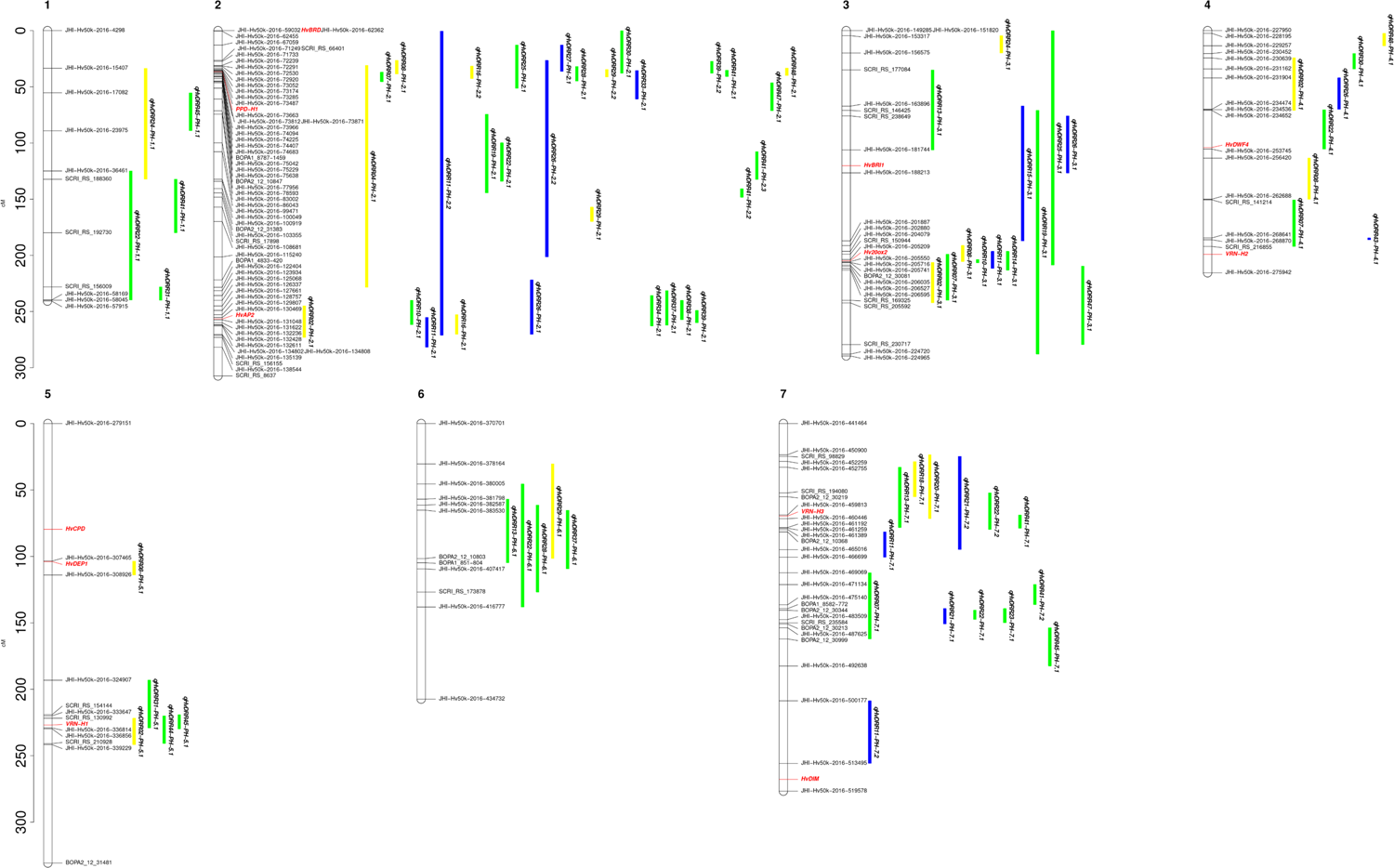
Genetic position of the QTL detected in single population analysis for plant height projected to the consensus map. The position of the QTL confidence intervals is shown as a vertical bar to the right of the chromosome. The color of the bar indicates if the sub-population was obtained by crossing two landraces (yellow), two cultivars (blue), or a landrace and a cultivar (green). The known regulatory genes previously described to be responsible for plant height regulation and their genetic position are reported in red. The positions of the markers at the borders of each QTL are also reported.

Out of 89 QTL identified in the single population analysis for FT, 43 mapped to chromosome 2H (Figure 5). A cluster comprising 21 QTL was located at the beginning of chromosome 2H. The region covered by the confidence interval of these QTL included *Ppd-H1*. Also for other major effect genes, responsible for the control of FT, QTL clusters had been identified: six QTL at the end of chromosome 4H, whose confidence intervals included *Vrn-H2*, ten QTL at the beginning of chromosome 5H, a region in which *Vrn-H1* was positioned, and eleven QTL at the beginning of chromosome 7H in which *Vrn-H3* was located. Other QTL comprised additional genes within their confidence intervals such as *HvELF3*, *HvCEN*, *Hv20ox2* (*sdw1/denso)*, *HvFT4* (Pieper *et al*., 2021), and *HvAP2* (Supplementary Table 5). Single population analysis for PH identified 80 QTL, where these QTL were characterized by wider confidence intervals compared to those detected for FT (Figure 6, Supplementary Table 6). As observed for FT, the chromosome with the highest number of QTL was 2H. A cluster, including 14 QTL, comprised within their confidence interval *Ppd-H1*. Other clusters of QTL that included in their confidence interval *HvAP2*, *Hv20ox2* (*sdw1/denso*), *Vrn-H1*, and *Vrn-H3* had been identified (Supplementary Table 6).

However, we identified 16 QTL for FT and 31 QTL for PH for which no genes previously described for the control of the trait were included within their confidence interval (Supplementary Tables 5-6). Among the 16 QTL detected for FT, the QTL with the lowest number of genes in the confidence interval were *qHvDRR02-FT-5.1* and *qHvDRR31-FT-5.2.* The QTL comprised 52 and 71 genes, respectively, which reduced to 35 and 45 when neglecting the low confidence genes. *qHvDRR31-FT-5.2* was with 3.4 cM the QTL with the shortest genetic confidence interval. For PH, *qHvDRR48-PH-*4.1 was the QTL with the lowest number of genes in its confidence interval (115 low and high confidence or 79 high confidence genes). The QTL with the shortest confidence interval was *qHvDRR22-PH-7.1* with 3.9 cM.

Eight sub-populations showed significant epistatic interactions between loci on a genome-wide scale. In total, ten significant epistatic interactions were detected. Nine interactions were detected for PH and one for FT. Two epistatic interactions each were observed for populations HvDRR34 and HvDRR44 (Supplementary Table 8).

### 3.5 Allele mining

For FT, 21 sub-populations showed a QTL that included *Ppd-H1*. For 14 of these, a QTL that included *Ppd-H1* was also identified for PH. A total of 16 of the 21 sub-populations were polymorphic for the causal SNP 22 of *Ppd-H1* (Turner *et al*., 2005). However, five sub-populations (HvDRR02, HvDRR04, HvDRR20, HvDRR23, and HvDRR48), for which the QTL confidence intervals comprised *Ppd-H1*, did not segregate for this polymorphism. All non-polymorphic sub-populations for SNP 22 had HOR1842 or IG128104 as parental inbred lines (Supplementary Table 1). Through Sanger sequencing, we identified the presence of a unique SNP in HOR1842 and IG128104 in the CCT domain of *Ppd-H1* (Figure 7). The primers used to amplify *Ppd-H1* are listed in Supplementary Table 9. Based on the SNP position on the *Ppd-H1* coding sequence of Morex, we refer to it as SNP 1945. SNP 1945 determines the synthesis of a threonine instead of an alanine (Supplementary Figure 4). We then used the SIFT algorithm to predict the effect of this SNP on the phenotype. The effect of this polymorphism was classified by the SIFT algorithm as deleterious.

**Figure 7:**
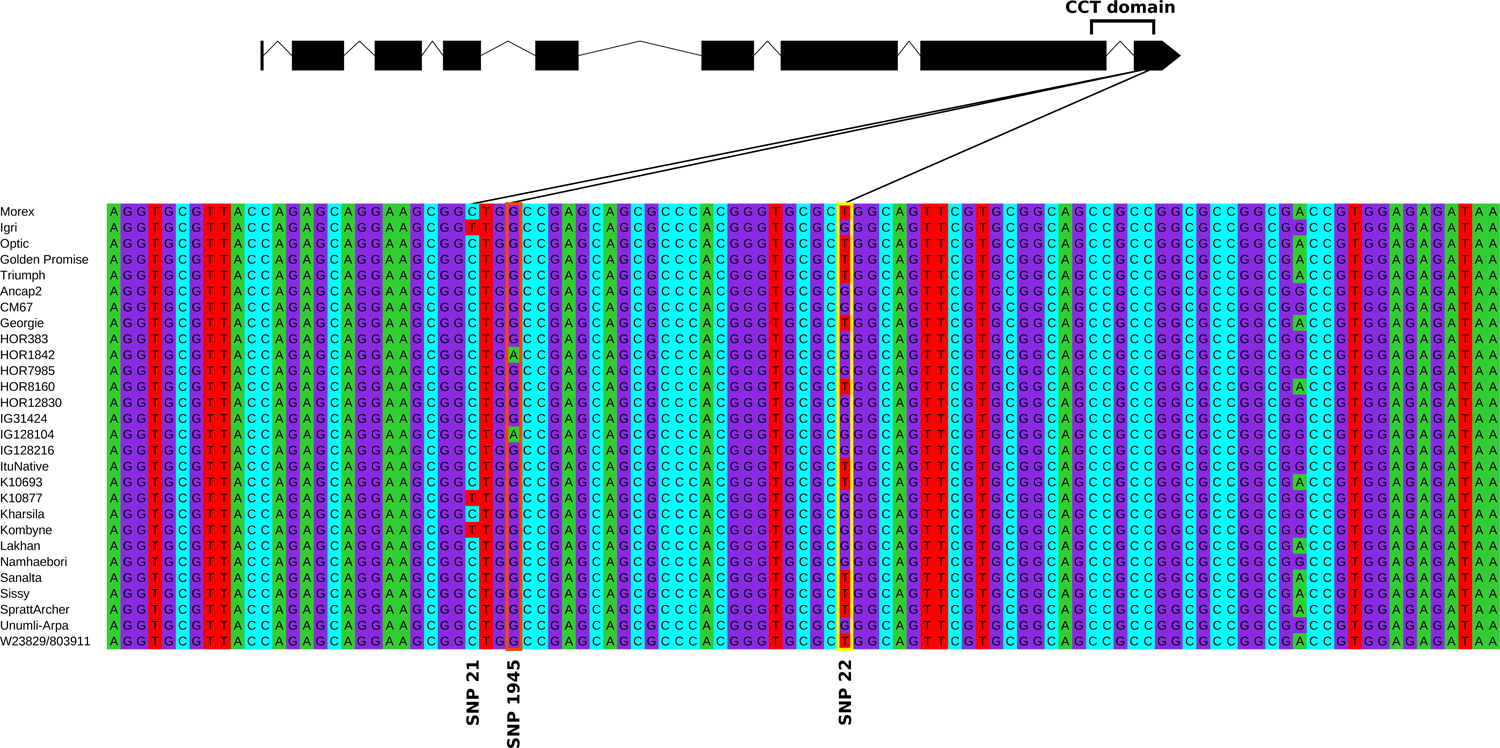
Genomic sequence of the last exon of *Ppd-H1* of Morex, Igri, Optic, Golden Promise, Triumph, and the 23 parental inbreds of the HvDRR population. SNP 22 is highlighted in yellow, SNP 1945 in orange. On top, the gene structure of *Ppd-H1* is given. Lines indicate the positions of SNP 21, SNP 22 (Turner *et al*, 2005), and SNP 1945 within the last exon.

At the *Vrn-H2* locus, an FT QTL was detected in six sub-populations. We evaluated by PCR, as described in Karsai *et al*. (2005) (Supplementary Table 9), the presence/absence of the three causal *Vrn-H2* genes. Out of six sub-populations, five were polymorphic for the genes regulating the *Vrn-H2* locus. In HvDRR29, both parental inbred lines, HOR8160 and IG128216, had the complete set of genes (Supplementary Figure 5).

### 3.6 Candidate gene analysis

The candidate gene analysis was performed for the QTL detected in the SP analysis that did not carry in their confidence interval previously reported genes controlling the trait under consideration, explaining ≥ 15% of the phenotypic variance, and having a confidence interval ≤ 30 cM. For these QTL, we combined the results of QTL mapping with variant calling data and results from WGCNA. Through WGCNA, 27 different gene modules were detected across all the expressed genes in the barley genome (Supplementary Figure 6). The correlation of the gene expression of modules and the adjusted entry means ranged from −0.52 to 0.49 for FT and from −0.54 to 0.47 for PH. Interestingly, the module with the highest correlation was the same for both traits. After selecting genes within the QTL range and which were included in one of the three modules with the highest or the lowest correlation (Supplementary Figure 7), we searched for candidate genes. The most represented class of genes for the two traits was that of receptor-like kinase, followed by genes involved in the ethylene pathways, and genes coding for F-box proteins (Supplementary Table 10).

In addition to the functional based candidate gene analysis, we used association genetics in the diversity panel to fine map the selected QTL using the genome-wide genotyping-by-sequencing data of Milner *et al*. (2019). For FT, none of the polymorphisms in the QTL confidence intervals was significantly associated with the phenotype. For PH, we identified four significant SNPs associated with the phenotype (Supplementary Figure 8).

## 4 DISCUSSION

With this study, we aimed to obtain a comprehensive overview of the environmental and genotypic contributions to the regulation of flowering time and plant height in barley. We performed MPP and SP analysis to elucidate the genetic complexity underlying the control of FT and PH. Finally, we identified candidate genes and new allelic variants using additional approaches such as WGCNA and association genetics.

### 4.1 Flowering time variation is less environmental sensitive than that of plant height

We observed that relative to the genotypic variance, the variance components of environment and genotype by environment interaction were higher for PH than FT (Table 1). This finding was in discordance with a previous study where the variance component of the genotype-environment interaction was higher for FT than for PH (Rodriguez *et al*., 2008). This result might be explained thereby by that the environments of our study differed mainly with respect to soil, precipitations, and temperature, which influence PH more strongly than FT (Li *et al*., 2003). In contrast, latitudinal differences, which heavily impact FT (Kikuchi and Handa, 2009), were very small among our environments. An additional explanation is the limited number of studied genotypes by Rodriguez *et al*. (2008), which reduced the precision to estimate variance components. Nevertheless, we observed for both traits high to very high heritabilities indicating that the adjusted entry means calculated for FT and PH are very suitable to unravel the genetics of both traits (Table 1).

### 4.2 The double round-robin population shows high variability of flowering time and plant height

We observed in our study, with a range of adjusted entry means of 51.2 to 105.7 days as well as 12.6 to 101.2 cm for FT and PH, respectively, a high phenotypic diversity among the RILs of the HvDRR population (Figure 1). This range was considerably higher than that observed in earlier studies (Afsharyan *et al*., 2020; Arifuzzaman *et al*., 2016; Cuesta-Marcos *et al*., 2008; Maurer *et al*., 2015; Nice *et al*., 2017). Also, the standard deviation of the adjusted entry means of the RILs of the HvDRR population was higher for both traits than that described in previous studies (Pauli *et al*., 2014; Wang *et al*., 2014) (Supplementary Table 3). These observations might be due to the higher number of RILs but also because of the selection of the 23 parental inbreds with maximal genotypic and phenotypic richness. In addition, the variation observed for FT and PH in the diversity panel of 224 spring barley inbreds (Pasam *et al*., 2012) was similar to that observed in individual sub-populations. However, it was considerably smaller (FT) and more influenced by a few outliers (PH) compared to the diversity observed in the entire HvDRR population (Figure 1). These findings of high phenotypic variability, combined with high heritability values and high-quality genotypic data suggest that the double-round-robin population is a very powerful tool for exploring and detecting new genetic variants underlying the control of agronomic traits in barley, also compared to association mapping panels.

### 4.3 QTL analyses uncovered the role of the genetic background in determining the correlation between FT and PH

We observed that the correlation between FT and PH was different among the HvDRR sub-populations (Figure 2; Supplementary Figure 2). These results are in agreement with those of previous studies where positive and negative correlations have been identified between FT and PH (Von Korff *et al*., 2006; Maurer *et al*., 2016; Nice *et al*., 2017; Schmalenbach *et al*., 2009), although only one direction of the correlation index, positive or negative, was observed in each of these studies. The high variability of the correlation values could be due to the great genotypic diversity of the parental inbreds used in our study. In order to understand this aspect better, we considered in detail the co-located QTL for FT and PH in the SP analysis.

Considering the sub-populations in which the adjusted entry means of the two traits had the most negative correlation (HvDRR10, HvDRR11, and HvDRR43), all QTL detected for FT were also detected for PH, although additional QTL were observed for the latter trait (Supplementary Tables 5-6). For four of the five FT/PH QTL pairs, the parental inbred line conferring a positive additive effect for PH revealed a negative additive effect for FT and vice versa (Supplementary Tables 5-6; Supplementary Figure 2). At the same time, the three sub-populations with the highest correlation coefficient between FT and PH (HvDRR19, HvDRR28, and HvDRR29) had QTL falling within the same interval for the two traits (Figures 5-6). In this case, however, for the three overlapping pairs of QTL, the positive additive effect was given by the same parental inbred for both traits (Supplementary Tables 5-6; Supplementary Figure 2).

In order to increase the resolution of the dissection of the genetic origin of the correlation between FT and PH, we exploited the MPP analysis. For each of the two studied traits, we identified 21 QTL through MPP analysis (Figure 3). The QTL profiles obtained through MPP analysis for both traits had peaks falling within neighboring regions for the main genes previously reported to control FT and PH, such as *Ppd-H1* and the three vernalization genes. The diversity of the parental inbreds and the large number of sub-populations as well as the total RILs resulted in a high mapping resolution that led to narrow confidence intervals. In our study, we observed a pleiotropic effect only for two QTL pairs (*FT-MP-Q3*/*PH-MP-Q3* and *FT-MP-Q19*/*PH-MP-Q20*). *FT-MP-Q3*/*PH-MP-Q3*, included in their common interval HORVU.MOREX.r3.1HG0075860 which was functionally annotated as transcription factor, while *FT-MP-Q19*/*PH-MP-Q20*, which explained a higher percentage of phenotypic variance compared to *FT-MP-Q3*/*PH-MP-Q3*, included in their intervals *Vrn-H3*. For all other QTL, the effect was separated by recombination.

Therewith, our results suggest that FT and PH variations are, with the exception of two QTL, caused by independent genetic factors (Figures 2-3-6-7; Supplementary Tables 4-5-6).

### 4.4 Multi-parent and single population analyses revealed new genome regions as well as genomic variants involved in the control of flowering time and plant height

The number of QTL identified through MPP analysis (Figure 3; Supplementary Table 4) was, with 21 each, higher compared to the maximum number of QTL, 13 for FT and 20 for PH, that were detected in earlier studies using bi- and multi-parental populations of spring barley (Arifuzzaman *et al*., 2014; Cuesta-Marcos *et al*., 2008; Von Korff *et al*., 2006; Maurer *et al*., 2015; Nice *et al*., 2017; Pauli *et al*., 2014; Rollins *et al*., 2013; Schmalenbach *et al*., 2009). The only exception was the study of Hemshrot *et al*. (2019) where a total of 23 QTL were identified for FT. However, in Hemshrot *et al*. (2019), QTL with the same genetic position were detected which reduced the number of QTL with different genetic positions to 13. The reasons for the higher number of QTL detected in our study compared to earlier studies were most probably the greater number of RILs and environments as well as the selection of very diverse parental inbreds (Weisweiler *et al*., 2019) which both increased the statistical power to detect QTL (Stich, 2009). Our observation suggested that the genetic complexity of FT and PH is higher than initially reported. This conclusion is furthermore supported by the observation that for both traits the percentage of explained variance by a genomic prediction model was about twice the value of the variance explained by the QTL detected in the MPP analysis (Supplementary Table 7). This result suggested that even with about 4000 RILs many small effect QTL remain undetected.

The difference between the percentage of variance explained by a genomic prediction model and the variance explained by the QTL detected in the MPP analysis was greater in the case of PH compared to FT (Supplementary Table 7). This observation suggests that PH is more strongly influenced by small (and undetected) effect QTL than FT. In addition, the total proportion of variance explained by the detected QTL was with 25.7% lower for PH than for FT (37.4%). This trend was in agreement with the observation that epistatic interaction played a bigger role for PH than for FT (Supplementary Table 8).

The SP QTL analyses detected 89 QTL for FT and 80 for PH (Figures 5-6; Supplementary Tables 5-6). We determined the physical position of QTL reported in earlier studies (Afsharyan *et al*., 2020; Druka *et al*., 2011; Hemshrot *et al*., 2019; Laurie *et al*., 1994; Maurer *et al*., 2015; Nice *et al*., 2017; Pauli *et al*., 2014), wherever possible, and compared it to the QTL observed in our study. We observed for 166 QTL detected in our study a co-localization with earlier reported QTL. However, three QTL detected with SP analyses, one for FT and two for PH, did not overlap with other previously reported QTL. The novel QTL detected by SP were *qHvDRR30-FT-*3.1, *qHvDRR24-PH-3.1*, and *qHvDRR48-PH-4.1*. The percentage of variance explained by these QTL was relatively low for *qHvDRR30-FT-*3.1 (4.3%) but higher for *qHvDRR24-PH-3.1* (26.2%) and *qHvDRR48-PH-4.1* (19.5%). Because of the high percentage of variance explained by *qHvDRR24-PH-3.1*, we started fine mapping project of this QTL, as well as for *qHvDRR28-FT-2.2*, *qHvDRR41-FT-2.2, qHvDRR42-FT-3.1, qHvDRR22-PH-7.1, qHvDRR29-PH-2.1*, and *qHvDRR47-PH-2.1*.

We observed for 21 sub-populations an FT QTL whose confidence interval included the *Ppd-H1* locus. Five of these sub-populations (HvDRR02, HvDRR04, HvDRR20, HvDRR23, and HvDRR48) were not polymorphic for SNP 22 (Figure 7; Supplementary Table 1). SNP 22 is located within the CCT domain of *Ppd-H1*, one of the two main regulatory regions of the *Ppd-H1* causal gene (Turner *et al*., 2005). SNP 22 was described as the polymorphism responsible for the difference between the dominant and recessive allelic variant of *Ppd-H1* (Turner *et al*., 2005). Although more than 80 variants had been detected for *Ppd-H1* (Jones *et al*., 2008), SNP 22 was so far the only functionally characterized polymorphism for which a difference in phenotype has been reported. A further polymorphism of *Ppd-H1*, SNP 48 (Jones *et al*., 2008), had previously been associated with FT variation. However, the study of Sharma *et al*. (2020) did not observe hints that SNP 48 was the causal SNP of the *Ppd-H1* mutation. In addition, in none of the above mentioned five sub-populations, SNP 48 was segregating. All five sub-populations had HOR1842 or IG128104 as parental inbreds (Supplementary Table 1). From the whole genome sequencing data of the parental inbreds (Weisweiler *et al*., 2022), followed by Sanger sequencing, we identified a not previously reported polymorphism, SNP 1945, unique to HOR1842 and IG128104 (Figure 7). SNP 1945 is located within the CCT domain of *Ppd-H1* and it causes the synthesis of threonine instead of alanine (Supplementary Figure 4). This amino acid change was predicted by the SIFT algorithm as deleterious. In the sub-population HvDRR24, whose parental inbreds were HOR1842 and IG128104, we did not detect a QTL either for FT or for PH in the genome region of *Ppd-H1*. Together with the fact that HOR1842 and IG128104 originated from the same geographical region (south-central Asia), these observations support the hypothesis that HOR1842 and IG128104 might carry the same causal polymorphism. In addition, we observed that the additive effect for FT QTL co-locating with *Ppd-H1* was, with about 3.5 days, higher in sub-populations that segregated for SNP 22 compared to about 2.3 days for the five sub-populations that did not segregate for SNP 22 (Supplementary Table 5). For the latter sub-populations, the additive effect assumed a positive value for the RILs that inherited the *Ppd-H1* allele from HOR1842 or IG128104. These findings suggest that SNP 1945 is the causal polymorphism for the QTL in those sub-populations that are monomorphic for SNP 22 as well as a new functional allelic variant of *Ppd-H1*.

A similar observation was made for the QTL co-localizing with *Vrn-H2*. The *Vrn-H2* locus has been described as one of the main loci responsible for the difference between winter and spring barley varieties (Distelfeld *et al*., 2009). This difference is caused by the total deletion of a complex of three genes (*ZCCT-Ha, ZCCT-Hb*, and *ZCCT-Hc*) in the spring barley genotypes or, in the case of facultative genotypes, of a partial deletion (Fernández-Calleja *et al*., 2021; Karsai *et al*., 2005). Surprisingly, we observed for four of the HvDRR parental inbreds the complete set of *Vrn-H2* causal genes in spring varieties of barley (Supplementary Figure 5), which was previously reported only for winter varieties (Fernández-Calleja *et al*., 2021). This observation suggests that the plant response to the vernalization requirement may be more complex than previously assumed and not merely based on the presence/absence of the *Vrn-H2* genes. In addition, among the six sub-populations for which an FT QTL was detected at the *Vrn-H2* genome position, one sub-population, HvDRR29, was monomorphic for the number of *Vrn-H2* genes. Both parental lines, HOR8160 and IG128126 carried three *Vrn-H2* causal genes (Supplementary Figure 5). Similarly to *Ppd-H1*, it could be hypothesized that one of the parental lines of sub-population HvDRR29 carried a new functional allelic variant or that an additional gene, that acted on the phenotype in a similar way to *ZCCT-Ha:c*, was present within the same QTL confidence interval.

These two examples suggest that the genetic complexity of the studied traits might be higher than anticipated from the simple comparison of the list of co-localizing QTL and can now be resolved using multiple segregating populations together with next-generation sequencing of the parental inbreds. In addition, the cloning of the underlying genes will complement our understanding of the regulatory mechanisms of flowering time and plant height.

### 4.5 Candidate gene analysis for a subset of the QTL

We first extracted the polymorphic genes among the parental inbred lines within the confidence interval of the QTL that explained ≥ 15% of the phenotypic variance, had a confidence interval ≤ 30 cM, and did not carry in their confidence interval any previously reported gene controlling the trait under consideration. Then, we combined this screening with the results obtained from the WGCNA, selecting the three modules that showed each the lowest and highest correlation with FT and PH (Supplementary Figure 6).

Among the QTL detected for FT, *qHvDRR28-FT-2.2* was the one that had the highest percentage of explained variance and, at the same time, had the shortest genetic confidence interval. Two, out of the five candidate genes identified for this QTL, encoded for the pseudo-response regulator 3 (PRR3) HORVU.MOREX.r3.2HG0170150 and the ethylene-responsive transcription factor HORVU.MOREX.r3.2HG0170460 (Supplementary Table 10). Pseudo-response regulator is the same class of genes to which *Ppd-H1* belongs. The role of these genes is critical for the regulation of the plant circadian clock (Eriksson and Millar, 2003; Mizuno and Nakamichi, 2005) and it has been described, among the other functions, to be involved in the control of flowering time (Hayama and Coupland, 2004). Five different sub-groups belonging to this class of genes have been reported: PRR1, PRR3, PRR5, PRR7 (to which *Ppd-H1* belongs), and PRR9 (Matsushika *et al*., 2000). Phylogenetic analyses grouped the five sub-groups into three main clusters: PRR1, PRR5-PRR9, and PRR3-PRR7 (Nakamichi *et al*., 2020). Although genes belonging to all three clusters have been described to control flowering time or to be influenced by the photoperiod, the only cluster containing genes from grass species described to be dependent on the photoperiod and at the same time to control flowering time was PRR3-PRR7 (Nakamichi *et al*., 2020). Therewith this gene is an interesting target for further functional studies.

Genes responsible for ethylene biosynthesis are instead involved in a multitude of developmental processes throughout the plant life cycle, ranging from the early stages of plant development to the regulation of senescence (Bleecker and Kende, 2000). The concentration of ethylene also influences gene networks that regulate flowering in order to optimize the timing of the transition from the vegetative to the reproductive stage in relation to endogenous and external stimuli (Iqbal *et al*., 2017). Although further studies are needed to identify the pathways regulated by ethylene in barley, in rice, overexpression of an ethylene receptor (*ETR2*) was associated with delayed flowering (Hada *et al*., 2009). The delay was associated with an up-regulation of a homologous gene of GIGANTEA and TERMINAL FLOWER 1/CENTRORADIALIS (Hada *et al*., 2009), both of these classes of genes are involved in barley in the control of flowering since *HvGI* (Dunford *et al*., 2005) and *HvCEN* (Comadran *et al*., 2012) belong to them. Ethylene is also involved in plant growth (Dubois *et al*., 2018) and its role in vegetative development in plants has been described in barley (Patil *et al*., 2019). In addition to the one found in *qHvDRR28-FT-2.2*, we also identified two ethylene-responsive transcription factors (HORVU.MOREX.r3.7HG0685230 and HORVU.MOREX.r3.2HG0182430) in *qHvDRR22-PH-7.1* and *qHvDRR29-PH-2.1*. Besides being an ethylene-responsive transcription factor, HORVU.MOREX.r3.2HG0182430 also belongs to the same class of genes as *HvAP2*.

In addition to functional data, we used association genetics to fine map the detected QTL using the diversity panel which was evaluated in the same set of environments as the HvDRR population. For FT, none of the polymorphisms from Milner *et al*. (2019) that were located in the QTL confidence intervals were significantly associated (p < 0.05) with FT variation. The reason for this discrepancy was most probably that association mapping panels have a low power to detect marker-trait associations in the case of low frequency alleles (Myles *et al*., 2009), which is overcome by using segregating populations as in the HvDRR population. For PH, low significant marker-trait associations have been detected. However, only one of the polymorphisms was in proximity (< 150 kbp) to HORVU.MOREX.r3.3HG0222500, a candidate gene detected for *qHvDRR24-PH-3.1* using the WGCNA approach (Supplementary Table 10; Supplementary Figure 8). These results suggest that the integration of QTL analyses with other omics data sets supports the detection of candidate genes regulating traits of agronomic interest.

## 5 CONCLUSIONS

The great phenotypic variability observed for FT and PH in the HvDRR population suggests that this population will be a powerful genetic resource to detect new regulatory mechanisms that could allow to extend the barley cultivation area or its adaptation in changing environmental conditions. Furthermore, it was observed how environmental variables affected these traits and how the environmental component had a greater influence on plant height compared to flowering time. In addition, our study provides a comprehensive summary of the genetic architecture of FT and PH and serves as basis for future QTL cloning studies. Finally, the detection of novel QTL but also the observation that additional alleles or genes segregate at known loci like *Ppd-H1* and *Vrn-H2* suggests that the studied traits are genetically more complex than previously reported.

## 6 ACKNOWLEDGMENTS

We would like to thank our former colleagues George Alskief and Florian Esser for their technical support and for organizing and managing the field trials of the HvDRR population in the Cologne and Eifel location. We acknowledge Muenteha Yilmaz and Srinivasa Reddy Mothukuri for contribution to data collection and analyses. We thank the team of Saatzucht Breun for running the field trials in Quedlinburg. Computational infrastructure and support were provided by the Centre for Information and Media Technology at Heinrich Heine University, Düsseldorf.

## 7 AUTHORS CONTRIBUTIONS

Conceptualization: F.C., A.S. and B.S. Data analysis: F.C., A.S., D.V.I., F.Ca., P.W., M.W and B.S. Investigation: F.C and B.S. Resources: J.L., F.W. and B.S. Funding acquisition: B.S. Writing: F.C. and B.S.

## 10 DATA AVAILABILITY

The codes used for the calculation of the adjusted entry means, the single and multi-parent population QTL analyses, the epistatic QTL models, the WGCNA analysis, as well as the data sets of the adjusted entry means of the HvDRR population, the genetic haplotypes used to build the ancestral model, and the genotypic and phenotypic data used in the QTL analyses are deposited at https://github.com/cosenzaf/HvDRR_FT_PH. The data of membership of genes to gene modules used in the WGCNA are deposited in https://zenodo.org/record/7525604#.Y7_VgxXMLIW. Genetic maps and variant calling data can be obtained from Casale *et al*. (2022) and Weisweiler *et al*. (2022). Seeds of the RILs of the HvDRR population can be requested from the corresponding author.

## 12 TABLES AND FIGURES LEGENDS

**Supplementary Table 1:** Crossing scheme of the 45 HvDRR sub-populations. The name of the sub-populations is reported in the first column. In the second and third column are indicated the inbred lines that originated the sub-populations.

| Sub-population | Parent A | Parent B |
| --- | --- | --- |
| HvDRR02 | HOR1842 | IG31424 |
| HvDRR03 | Kharsila | IG31424 |
| HvDRR04 | HOR1842 | Kharsila |
| HvDRR07 | Sissy | HOR7985 |
| HvDRR08 | HOR7985 | W23829/803911 |
| HvDRR09 | Sissy | W23829/803911 |
| HvDRR10 | Sanalta | W23829/803911 |
| HvDRR11 | Sanalta | Sissy |
| HvDRR12 | SprattArcher | Sanalta |
| HvDRR13 | SprattArcher | HOR8160 |
| HvDRR14 | Georgie | HOR8160 |
| HvDRR15 | SprattArcher | Georgie |
| HvDRR16 | K10693 | HOR12830 |
| HvDRR17 | K10693 | Ancap2 |
| HvDRR18 | HOR383 | K10693 |
| HvDRR19 | HOR383 | Ancap2 |
| HvDRR20 | IG128104 | HOR383 |
| HvDRR21 | Ancap2 | Namhaebori |
| HvDRR22 | HOR383 | Namhaebori |
| HvDRR23 | IG128104 | Namhaebori |
| HvDRR24 | IG128104 | HOR1842 |
| HvDRR25 | W23829/803911 | Unumli-Arpa |
| HvDRR26 | Unumli-Arpa | Sanalta |
| HvDRR27 | Unumli-Arpa | SprattArcher |
| HvDRR28 | Unumli-Arpa | HOR8160 |
| HvDRR29 | HOR8160 | IG128216 |
| HvDRR30 | IG128216 | Georgie |
| HvDRR31 | Lakhan | IG128216 |
| HvDRR32 | IG128216 | K10877 |
| HvDRR33 | Georgie | Lakhan |
| HvDRR34 | K10877 | Lakhan |
| HvDRR35 | Lakhan | CM67 |
| HvDRR36 | HOR12830 | K10877 |
| HvDRR37 | K10877 | CM67 |
| HvDRR38 | HOR12830 | CM67 |
| HvDRR39 | HOR12830 | ItuNative |
| HvDRR40 | ItuNative | CM67 |
| HvDRR41 | ItuNative | K10693 |
| HvDRR42 | Ancap2 | ItuNative |
| HvDRR43 | Kombyne | Namhaebori |
| HvDRR44 | IG128104 | Kombyne |
| HvDRR45 | HOR1842 | Kombyne |
| HvDRR46 | Kharsila | Kombyne |
| HvDRR47 | IG31424 | Sissy |
| HvDRR48 | HOR1842 | HOR7985 |

**Supplementary Table 2:**
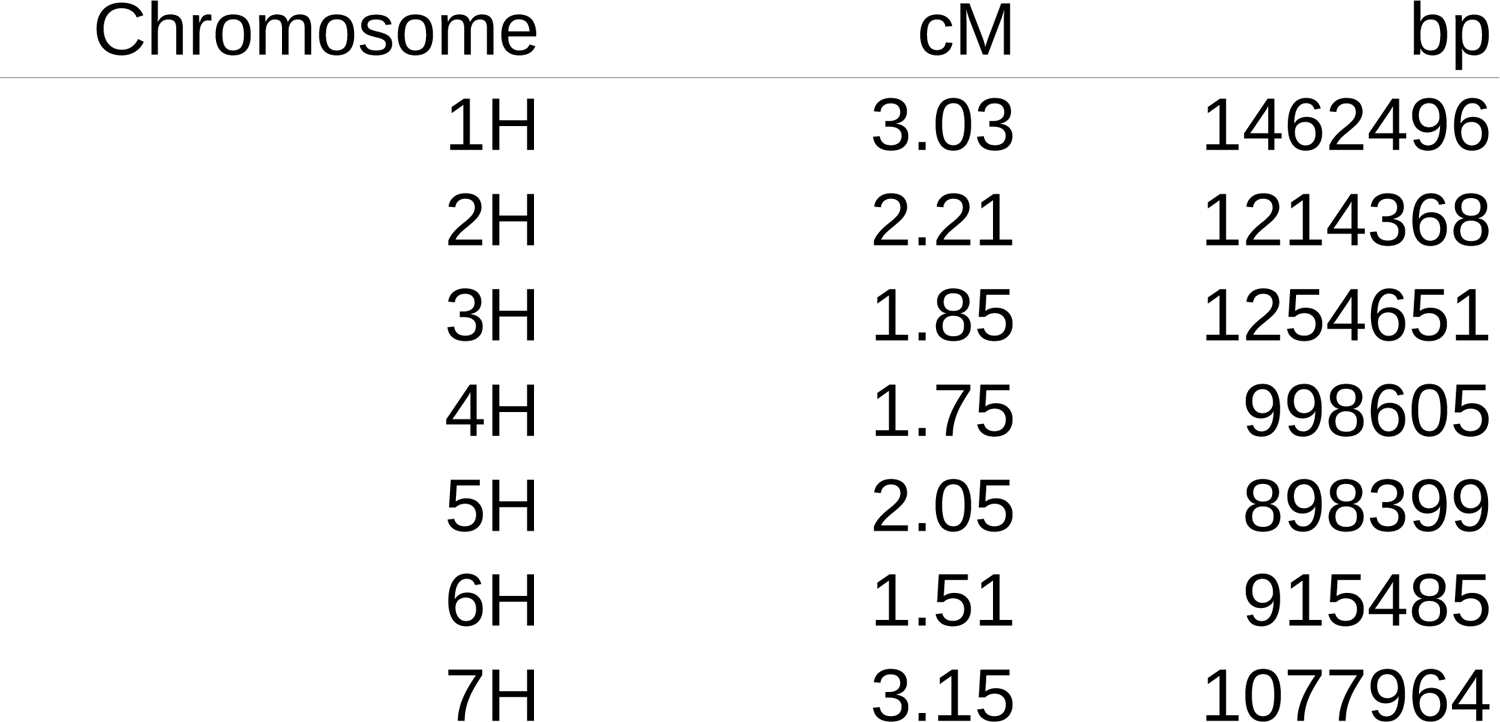
Genetic and physical distances for which the linkage disequilibrium measured *r²* reached a value of 0.2.

**Supplementary Table 3:** Average of the adjusted entry means, standard deviations (SD), and coefficients of variation (CoV) across all 45 sub-populations for flowering time, in days after sowing, and plant height, in cm.

| Sub-population | Flowering time (DAS) |  |  | Plant height (cm) |  |  |
| --- | --- | --- | --- | --- | --- | --- |
|  | Mean | SD | CoV | Mean | SD | CoV |
| HvDRR02 | 75.42 | 8.61 | 11.41 | 59.57 | 9.45 | 15.86 |
| HvDRR03 | 73.61 | 6.57 | 8.93 | 59.87 | 11.16 | 18.65 |
| HvDRR04 | 71.59 | 5.87 | 8.20 | 60.74 | 5.72 | 9.42 |
| HvDRR07 | 66.82 | 4.44 | 6.65 | 66.51 | 5.66 | 8.51 |
| HvDRR08 | 68.19 | 4.36 | 6.39 | 67.38 | 6.32 | 9.37 |
| HvDRR09 | 72.81 | 1.99 | 2.73 | 70.02 | 4.28 | 6.11 |
| HvDRR10 | 74.27 | 2.81 | 3.78 | 79.62 | 9.72 | 12.20 |
| HvDRR11 | 72.31 | 3.07 | 4.24 | 75.07 | 8.42 | 11.21 |
| HvDRR12 | 77.52 | 2.72 | 3.51 | 83.64 | 8.29 | 9.91 |
| HvDRR13 | 73.32 | 4.45 | 6.07 | 76.84 | 4.31 | 5.61 |
| HvDRR14 | 70.32 | 2.79 | 3.96 | 68.18 | 4.60 | 6.74 |
| HvDRR15 | 77.08 | 3.07 | 3.98 | 73.66 | 2.82 | 3.83 |
| HvDRR16 | 66.33 | 5.89 | 8.88 | 64.06 | 7.02 | 10.95 |
| HvDRR17 | 66.62 | 6.24 | 9.36 | 72.10 | 6.37 | 8.84 |
| HvDRR18 | 69.34 | 7.20 | 10.39 | 74.67 | 5.90 | 7.90 |
| HvDRR19 | 70.12 | 3.91 | 5.57 | 72.75 | 4.27 | 5.87 |
| HvDRR20 | 66.89 | 4.44 | 6.64 | 66.44 | 7.17 | 10.80 |
| HvDRR21 | 65.32 | 4.14 | 6.33 | 64.37 | 8.32 | 12.92 |
| HvDRR22 | 64.71 | 5.11 | 7.89 | 66.50 | 8.82 | 13.26 |
| HvDRR23 | 64.69 | 5.68 | 8.78 | 60.31 | 7.47 | 12.38 |
| HvDRR24 | 75.11 | 4.13 | 5.50 | 62.45 | 7.78 | 12.45 |
| HvDRR25 | 69.32 | 5.42 | 7.82 | 67.68 | 6.28 | 9.28 |
| HvDRR26 | 67.30 | 5.87 | 8.72 | 72.20 | 9.23 | 12.78 |
| HvDRR27 | 71.36 | 6.77 | 9.49 | 69.71 | 5.55 | 7.96 |
| HvDRR28 | 69.08 | 4.87 | 7.04 | 65.94 | 5.23 | 7.93 |
| HvDRR29 | 62.49 | 3.79 | 6.07 | 68.28 | 7.20 | 10.55 |
| HvDRR30 | 64.81 | 5.27 | 8.13 | 65.31 | 6.75 | 10.33 |
| HvDRR31 | 59.43 | 3.71 | 6.24 | 61.09 | 6.35 | 10.39 |
| HvDRR32 | 63.93 | 7.26 | 11.36 | 62.89 | 8.78 | 13.96 |
| HvDRR33 | 63.25 | 4.64 | 7.33 | 62.17 | 5.43 | 8.73 |
| HvDRR34 | 64.78 | 5.79 | 8.94 | 61.29 | 10.01 | 16.33 |
| HvDRR35 | 58.09 | 2.00 | 3.44 | 55.69 | 4.58 | 8.23 |
| HvDRR36 | 74.31 | 4.30 | 5.79 | 53.87 | 5.40 | 10.03 |
| HvDRR37 | 65.08 | 5.25 | 8.06 | 57.15 | 6.90 | 12.07 |
| HvDRR38 | 67.04 | 4.68 | 6.98 | 52.11 | 8.83 | 16.94 |
| HvDRR39 | 68.62 | 5.80 | 8.46 | 61.73 | 8.12 | 13.15 |
| HvDRR40 | 59.23 | 3.32 | 5.60 | 48.10 | 5.85 | 12.17 |
| HvDRR41 | 60.76 | 4.03 | 6.63 | 64.73 | 9.18 | 14.18 |
| HvDRR42 | 69.42 | 6.91 | 9.96 | 65.06 | 6.51 | 10.01 |
| HvDRR43 | 74.19 | 17.29 | 23.31 | 52.10 | 11.48 | 22.03 |
| HvDRR44 | 71.14 | 6.25 | 8.79 | 57.96 | 11.51 | 19.86 |
| HvDRR45 | 75.79 | 8.50 | 11.22 | 60.96 | 8.47 | 13.89 |
| HvDRR46 | 79.35 | 8.21 | 10.34 | 51.62 | 15.76 | 30.53 |
| HvDRR47 | 67.57 | 8.42 | 12.47 | 65.24 | 7.40 | 11.35 |
| HvDRR48 | 71.68 | 4.37 | 6.09 | 63.54 | 6.29 | 9.91 |
| Diversity panel | 69.63 | 5.12 | 7.35 | 67.38 | 9.44 | 14.01 |

**Supplementary Table 4:**
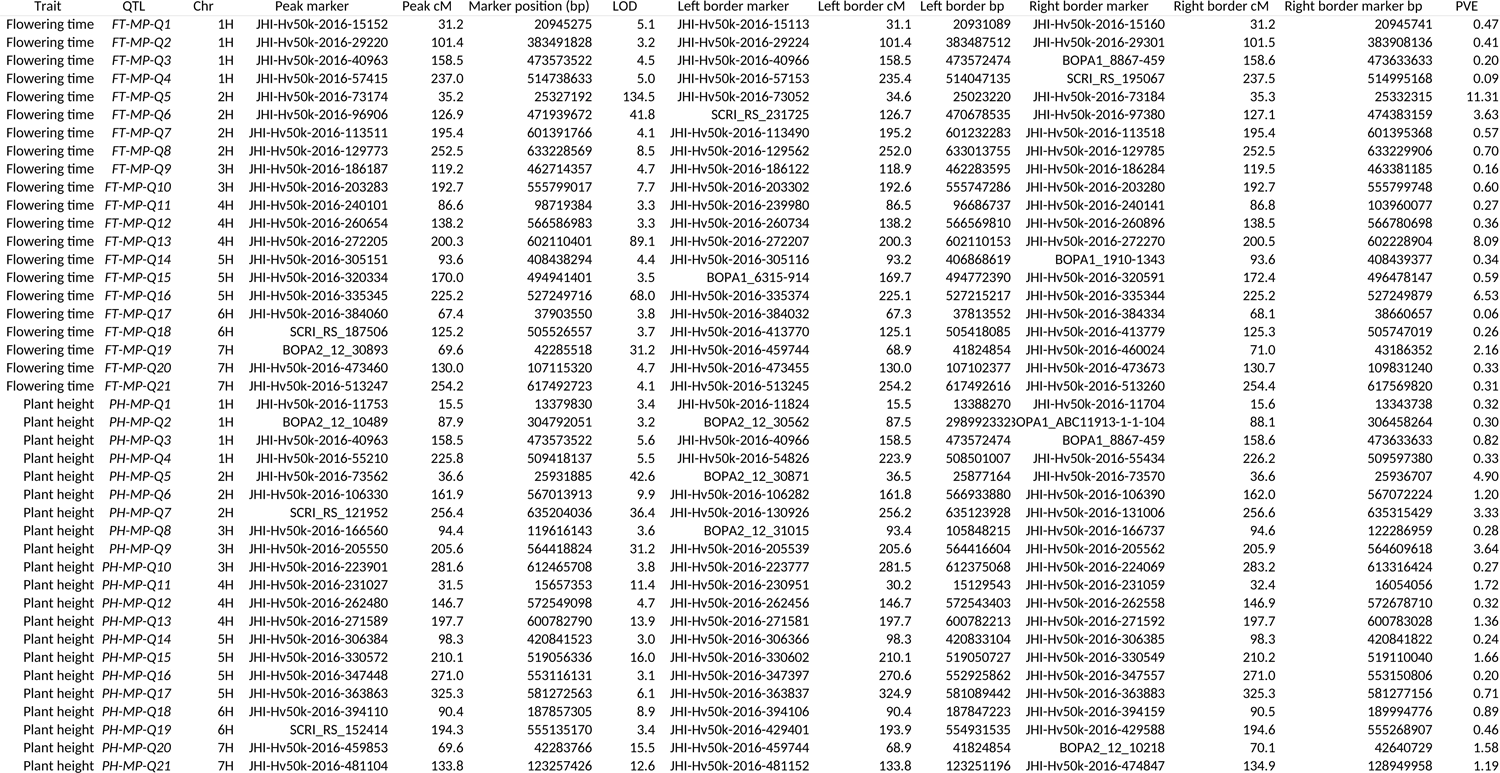
Summary of the results of the multi-parent population analysis for flowering time and plant height. Chr indicates the chromosome on which the QTL was detected, LOD the logarithm of odds, PVE the percentage of variance explained by the QTL.

**Supplementary Table 5:**
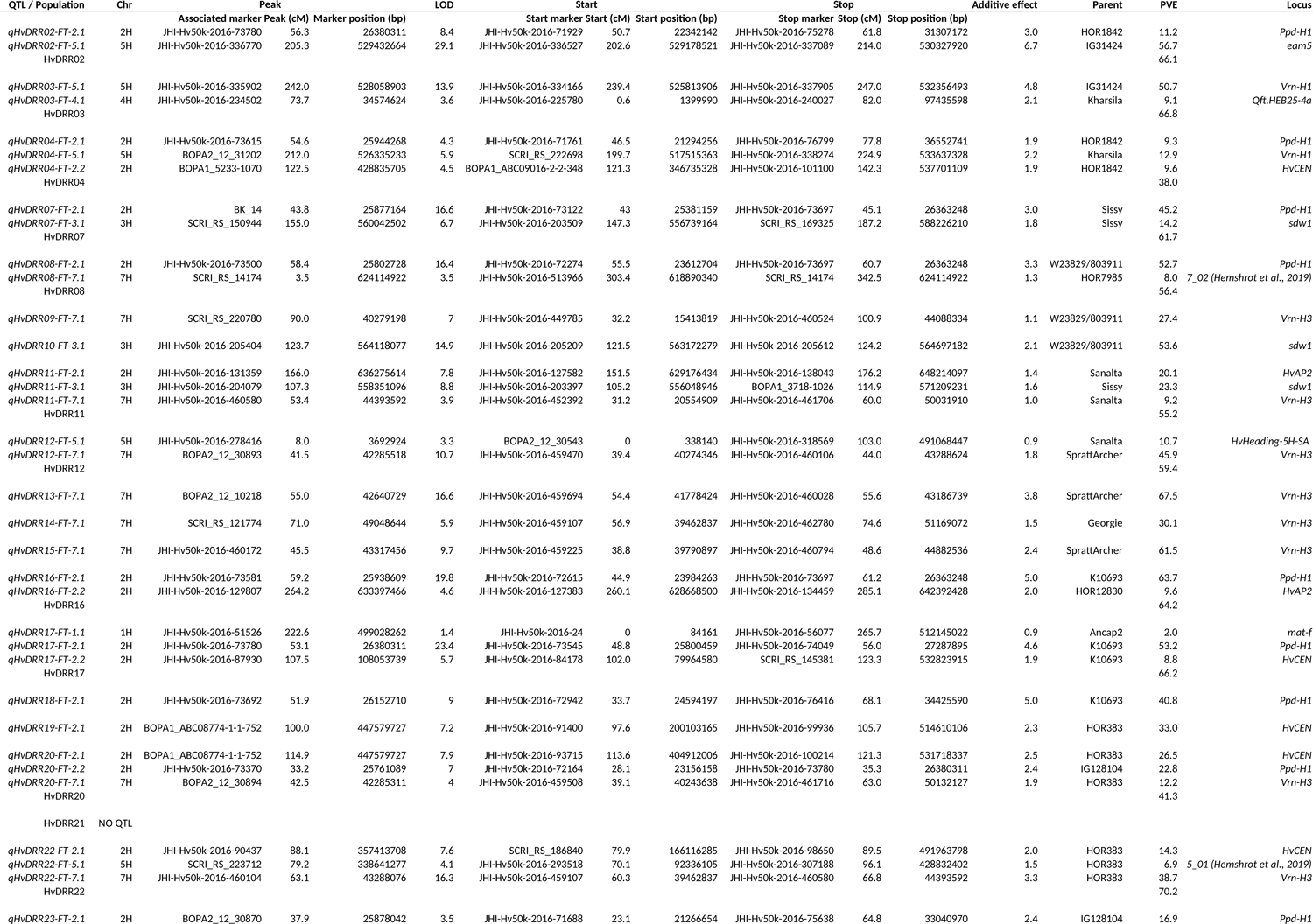
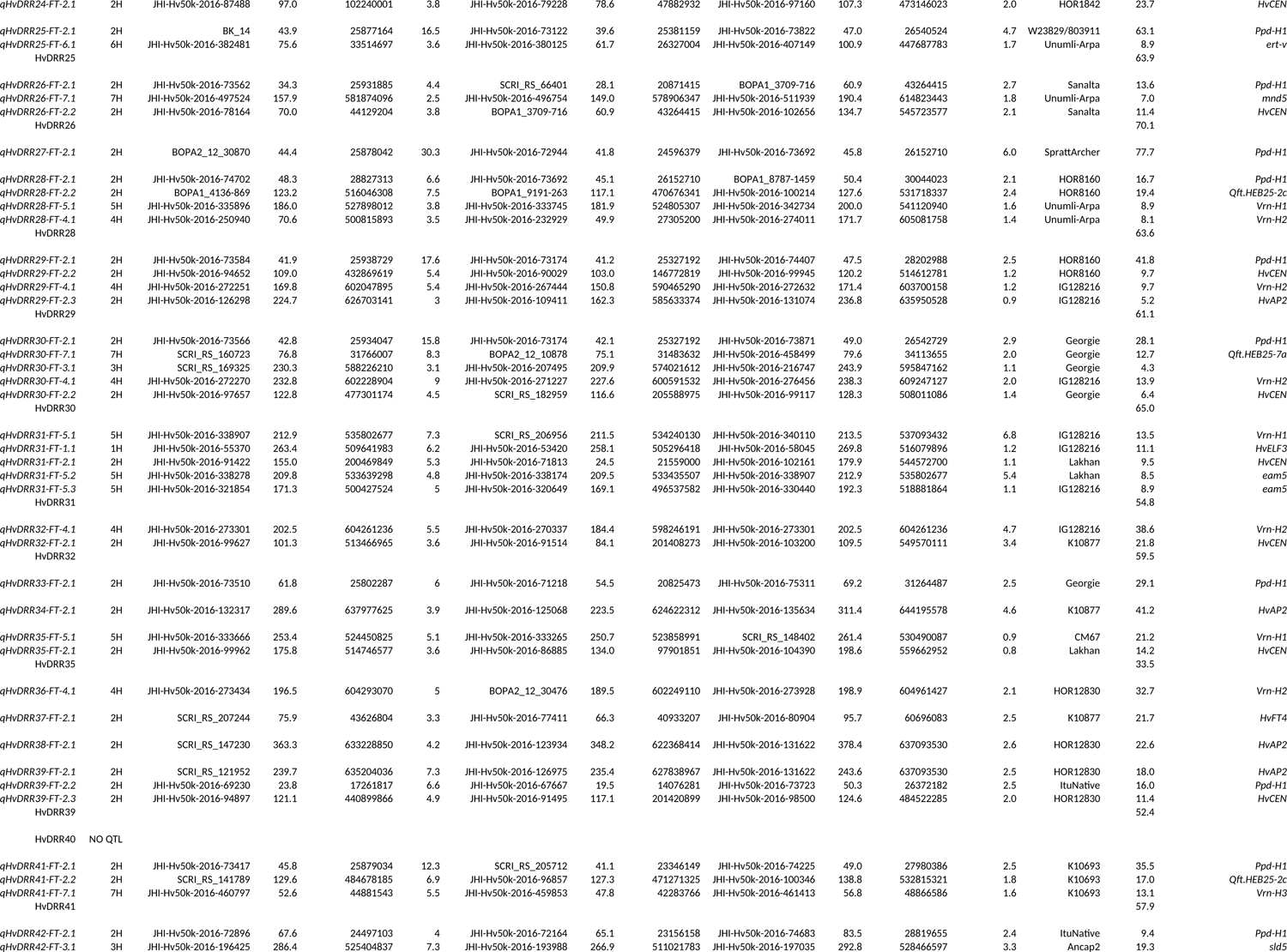
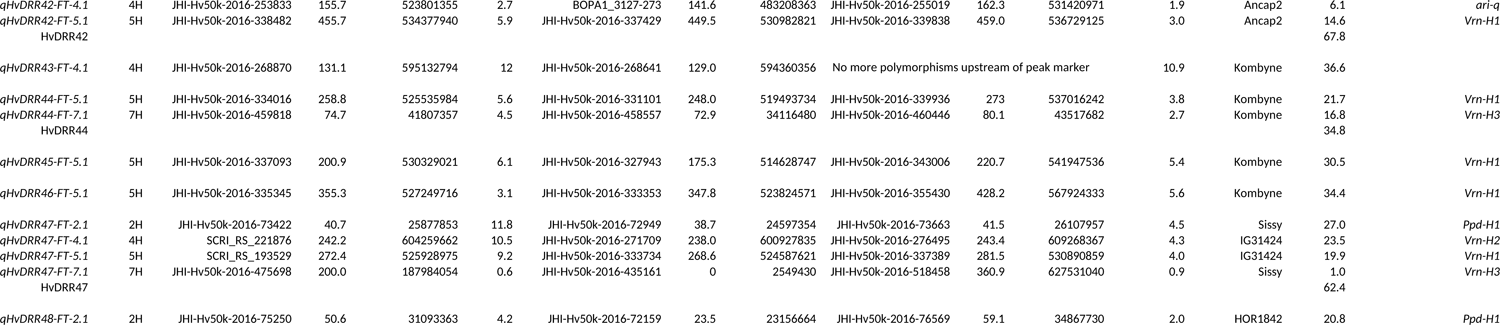
Summary of the results of the single population analysis for flowering time. The information regarding the peak and the borders of the confidence interval of each QTL are reported. Chr represents the chromosome on which the QTL was detected, LOD the logarithm of odds, and PVE the percentage of variance explained by the QTL individually and in a simultaneous fit. The additive effect is given in days after sowing.

**Supplementary Table 6:**
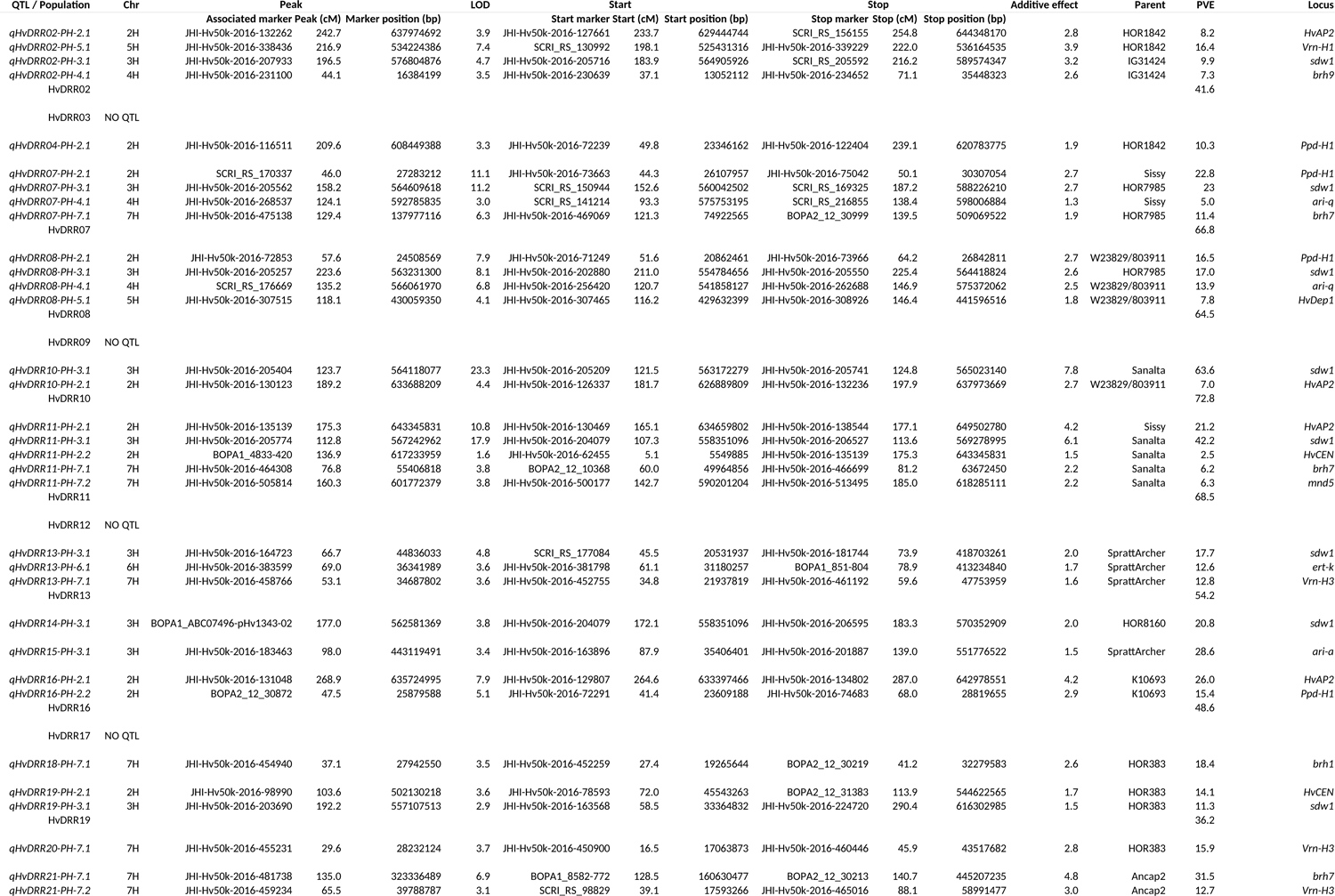
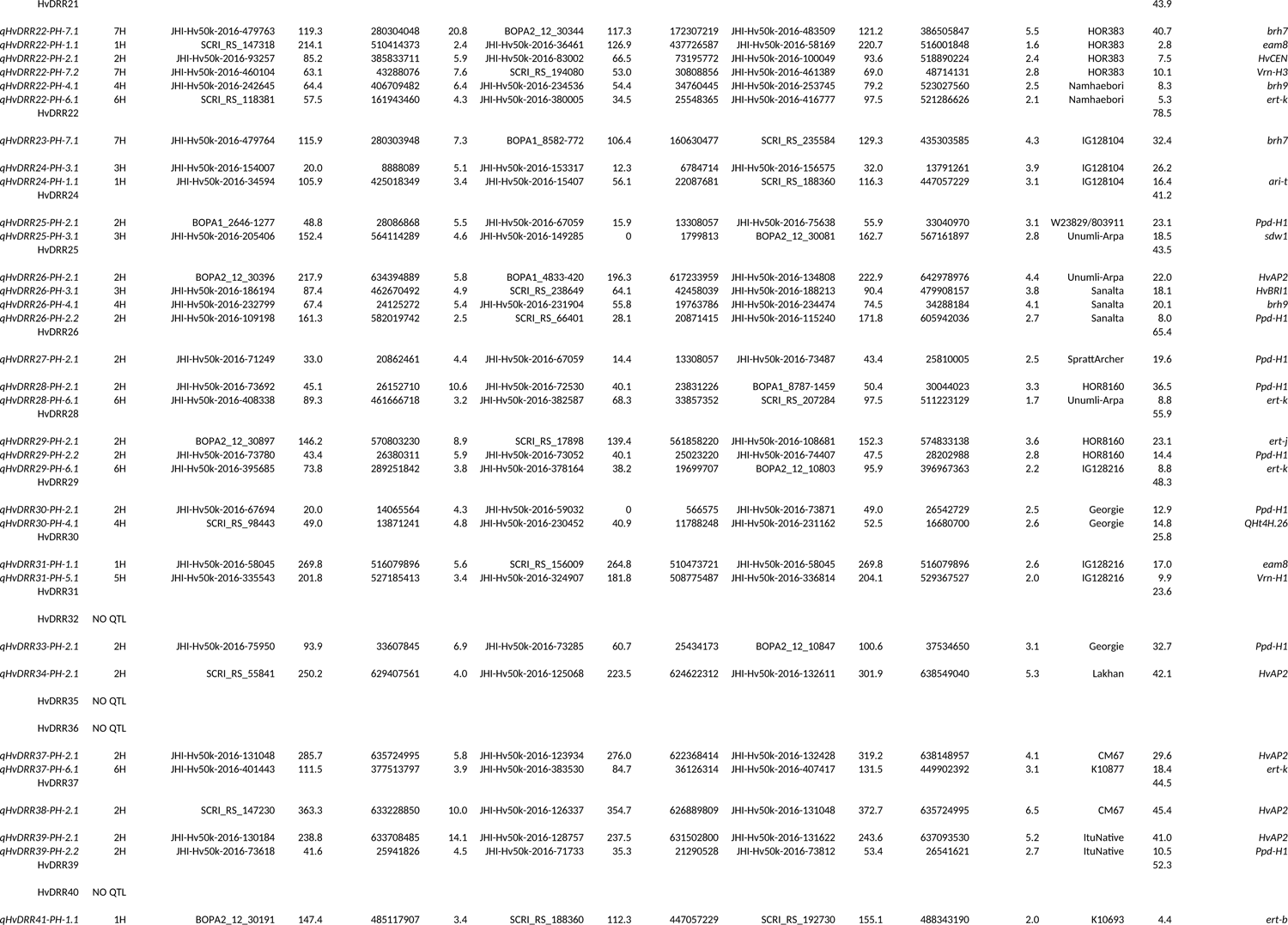
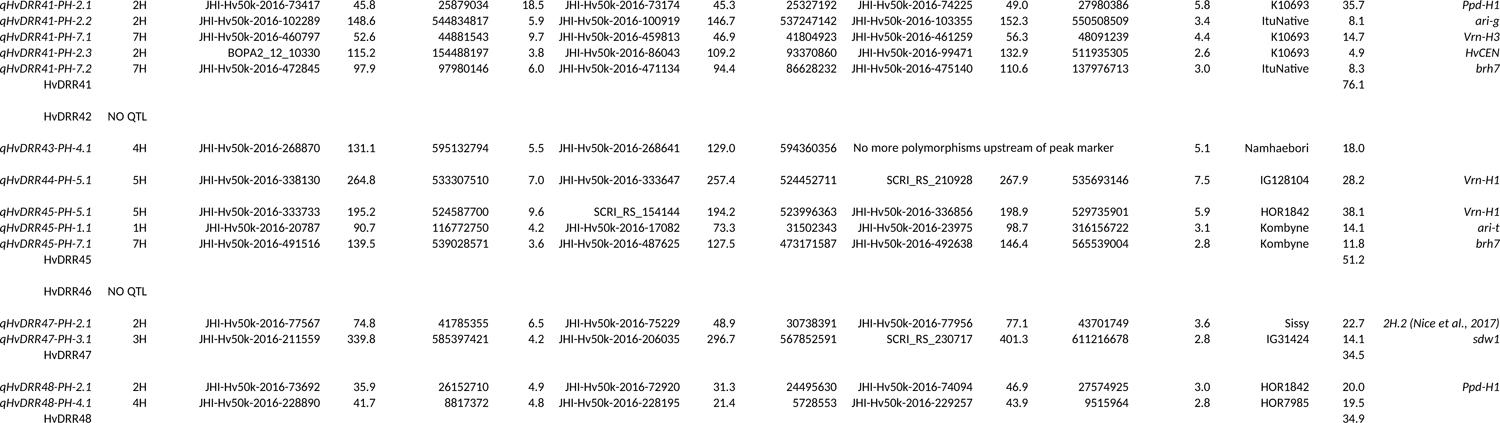
Summary of the results of the single population analysis for plant height. The information regarding the peak and the borders of the confidence interval of each QTL are reported. Chr represents the chromosome on which the QTL was detected, LOD the logarithm of odds, and PVE the percentage of variance explained by the QTL individually and in a simultaneous fit. The additive effect is given in cm.

**Supplementary Table 7:** Prediction ability of the genomic SNP marker data for flowering time and plant height without cross-validation (CV) and with five fold cross-validation across all sub-populations. SD indicates the standard deviation.

|  | Prediction ability across sub-populations |  | SD |
| --- | --- | --- | --- |
|  | Without CV | 5 Fold CV 20 runs | 5 Fold CV 20 runs |
| Flowering time | 0.89 | 0.766 | 0.015 |
| Plant height | 0.874 | 0.773 | 0.014 |

**Supplementary Table 8:** Genome-wide epistatic loci detected in the HvDRR population. LOD indicates the logarithm of odds of the interaction.

| Trait | Population | Chromosome locus 1 | Position locus 1 (cM) | Chromosome locus 2 | Position locus 2 (cM) | LOD | p-value |
| --- | --- | --- | --- | --- | --- | --- | --- |
| Flowering time | HvDRR13 | 5H | 107.5 | 6H | 87.5 | 6.74345 | 0.006 |
| Plant height | HvDRR02 | 5H | 205.0 | 7H | 72.5 | 11.5203 | 0 |
| Plant height | HvDRR09 | 1H | 172.5 | 7H | 97.5 | 4.273374 | 0.042 |
| Plant height | HvDRR17 | 1H | 2.5 | 1H | 7.5 | 4.125857 | 0.046 |
| Plant height | HvDRR34 | 4H | 165.0 | 5H | 180.0 | 4.260018 | 0.034 |
| Plant height | HvDRR34 | 4H | 160.0 | 7H | 215.0 | 4.484935 | 0.022 |
| Plant height | HvDRR41 | 1H | 122.5 | 5H | 225.0 | 3.985981 | 0.042 |
| Plant height | HvDRR44 | 2H | 75.0 | 5H | 262.5 | 11.46462 | 0.002 |
| Plant height | HvDRR44 | 5H | 262.5 | 7H | 195.0 | 11.14014 | 0.002 |
| Plant height | HvDRR45 | 1H | 85.0 | 5H | 197.5 | 5.781251 | 0.032 |

**Supplementary Table 9:**
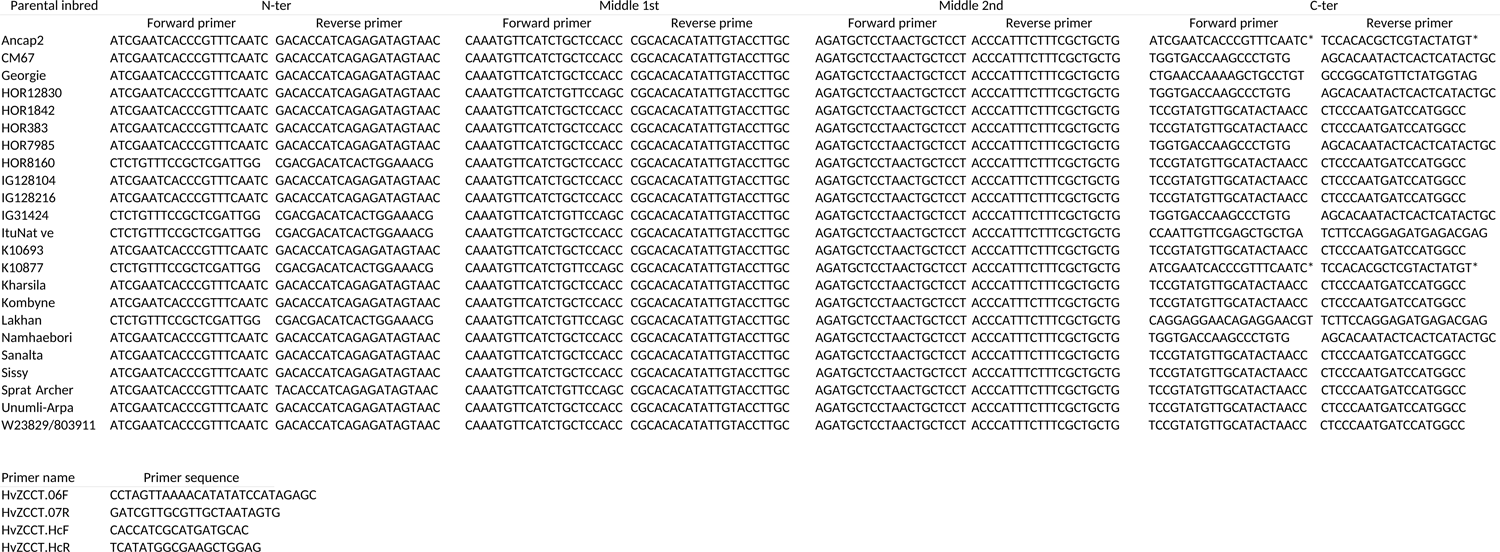
Lists of primers used to amplify *Ppd-H1* and *Vrn-H2*. The *Ppd-H1* primers are listed for each parental inbred. N-ter primers were used to amplify the start while the C-ter primers the end of the coding sequences. The primers pairs marked with * amplified the whole genet. Primers used to amplify *Vrn-H2* have the same nomenclature as described in Karsai *et al*. (2005).

**Supplementary Table 10:**
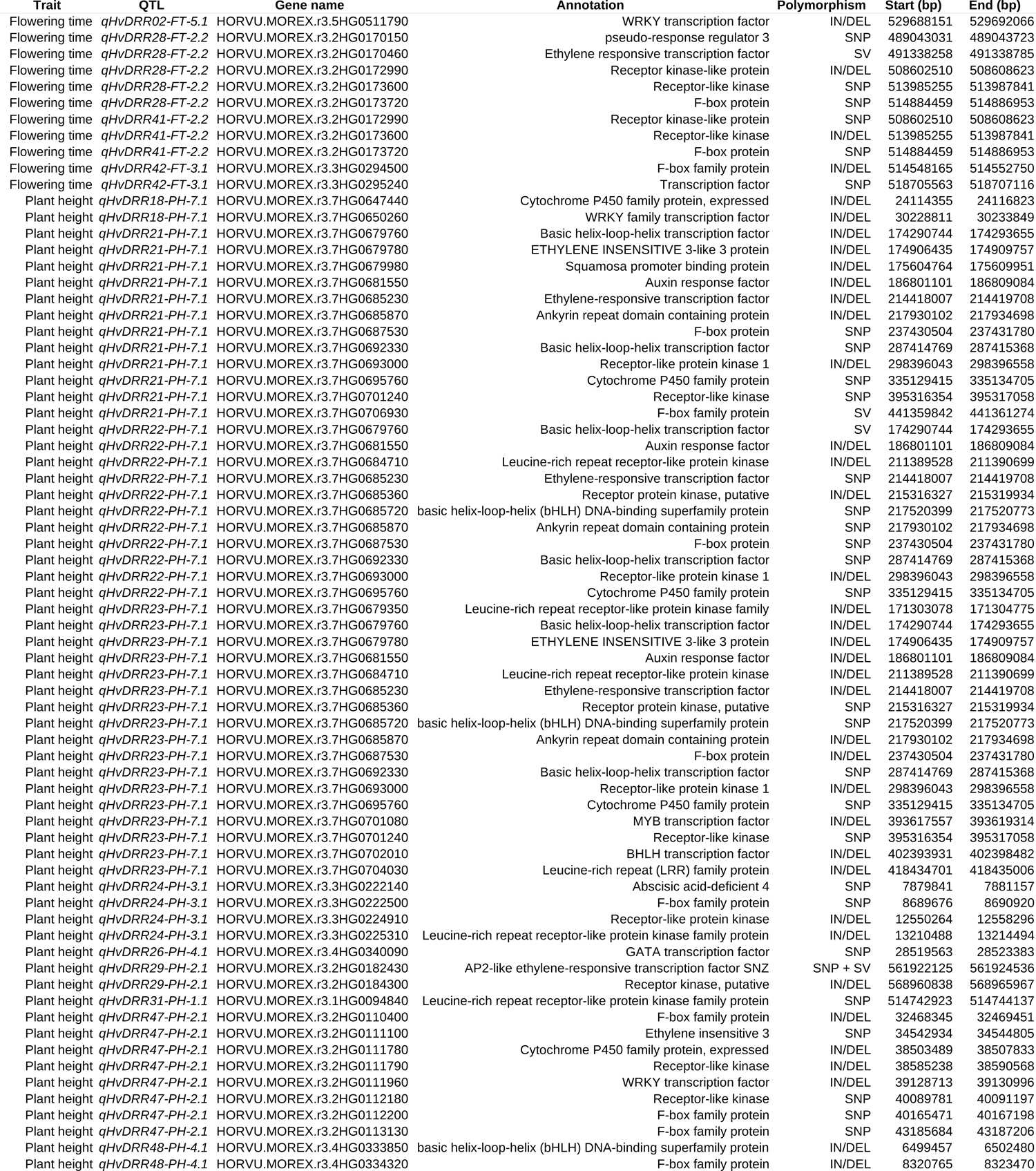
List of candidate genes in the confidence interval of selected QTL that carried a polymorphism among the parental lines. IN/DEL indicates an insertion or a deletion, SV indicates predicted structural variants.

---

**Supplementary Figure 1:**
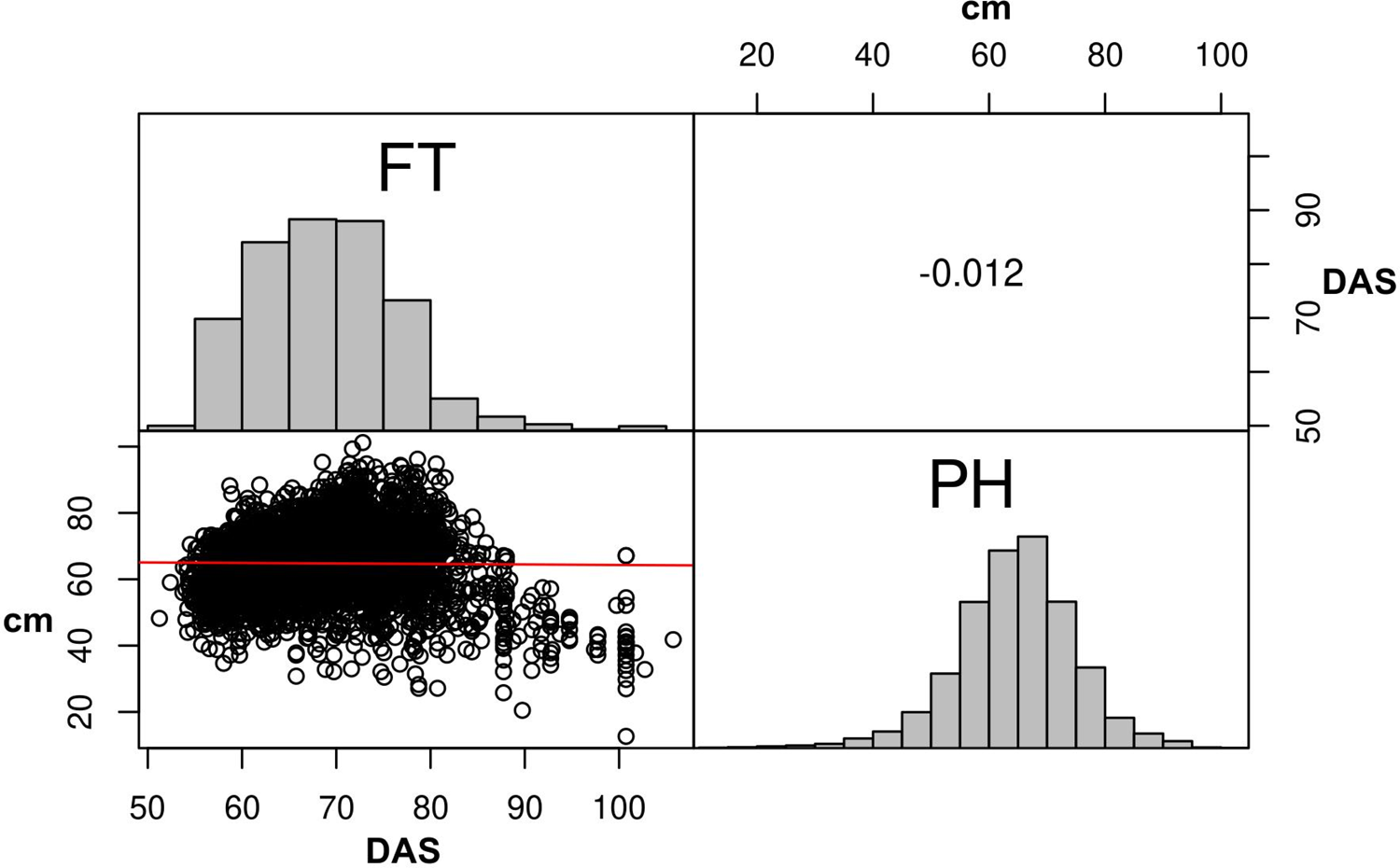
Histogram and correlation plot between flowering time (FT) and plant height (PH) across all 45 HvDRR sub-populations. Flowering time is reported in days after sowing (DAS) and plant height in cm.

**Supplementary Figure 2:**
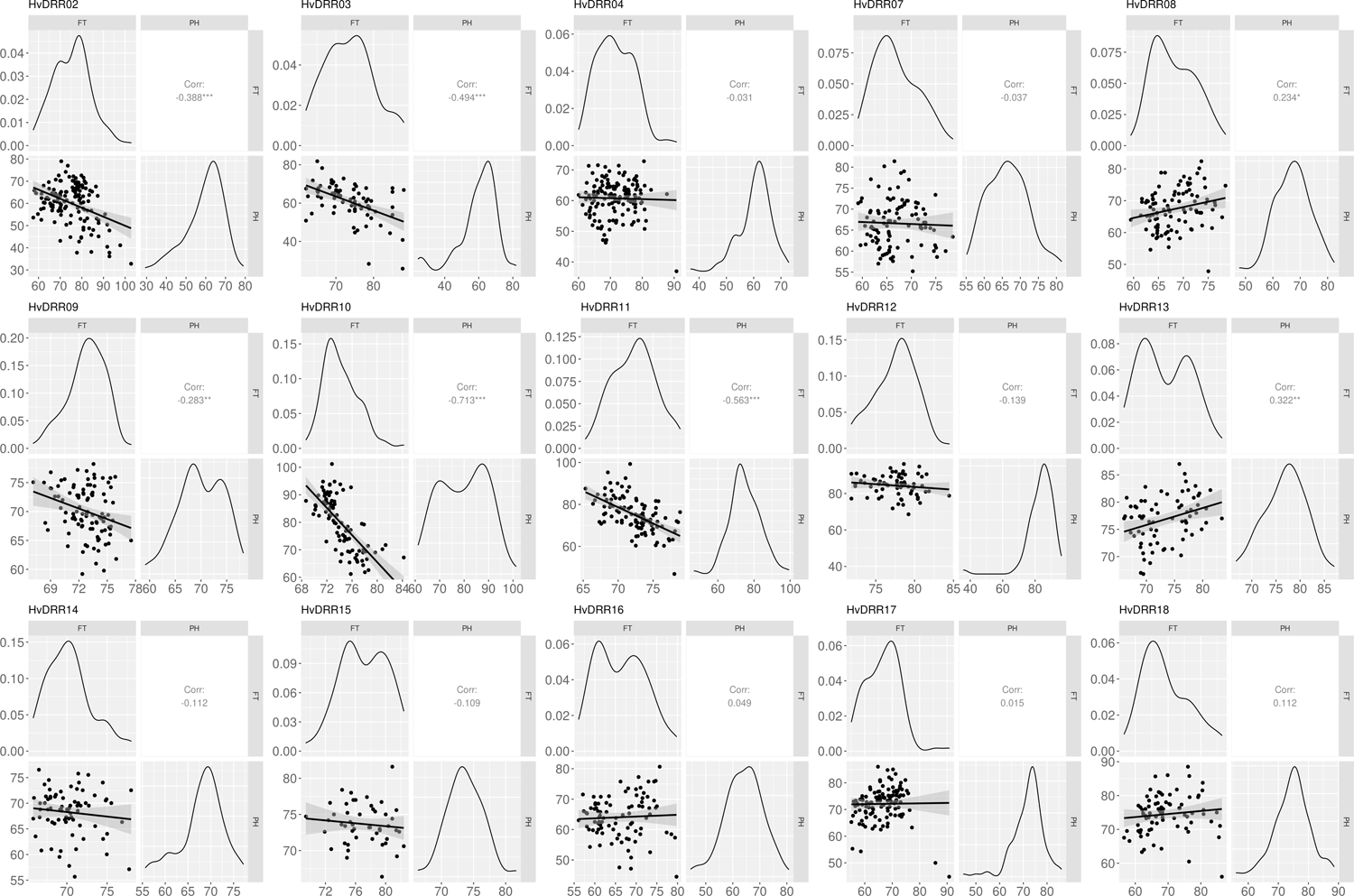
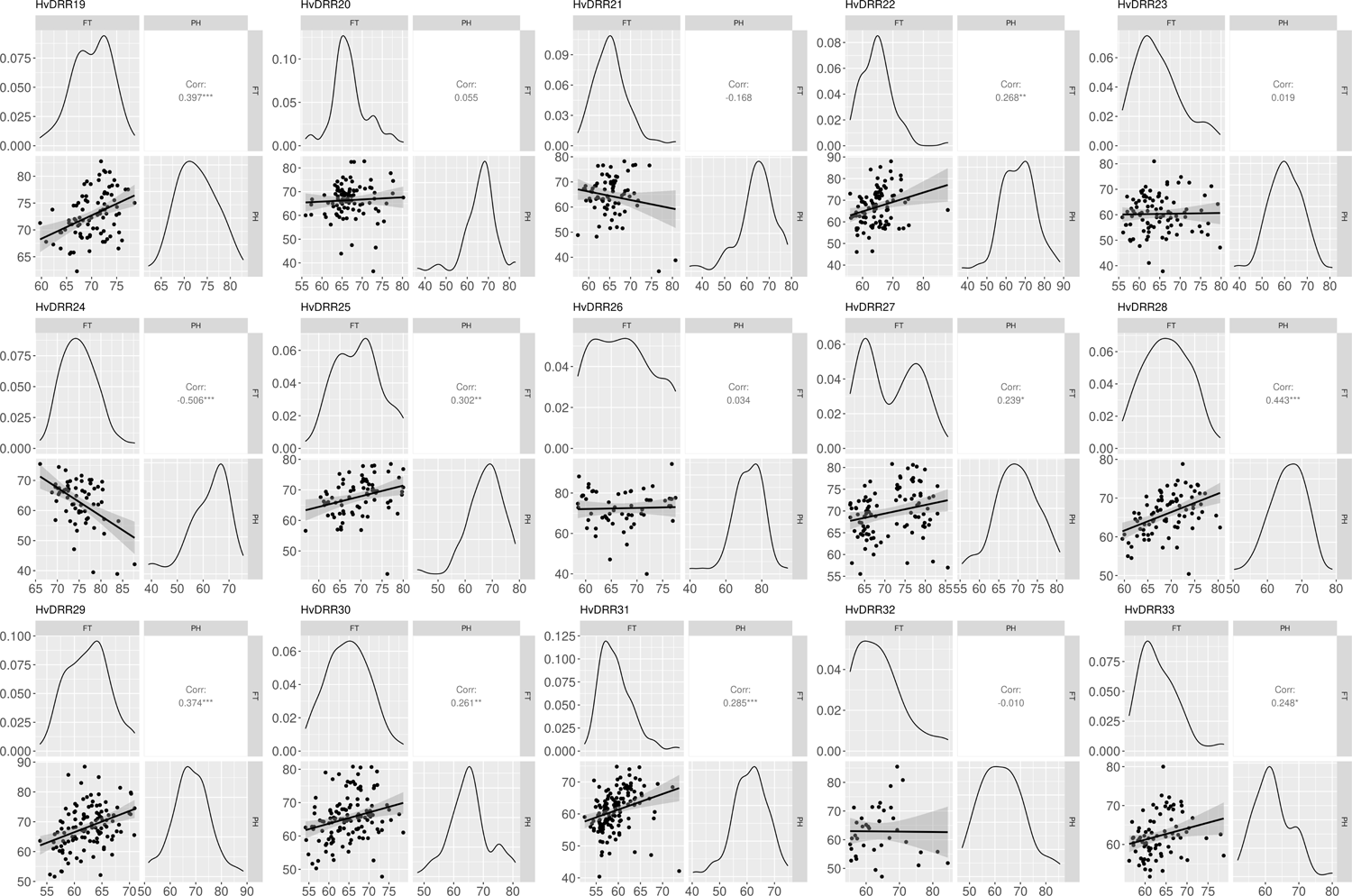
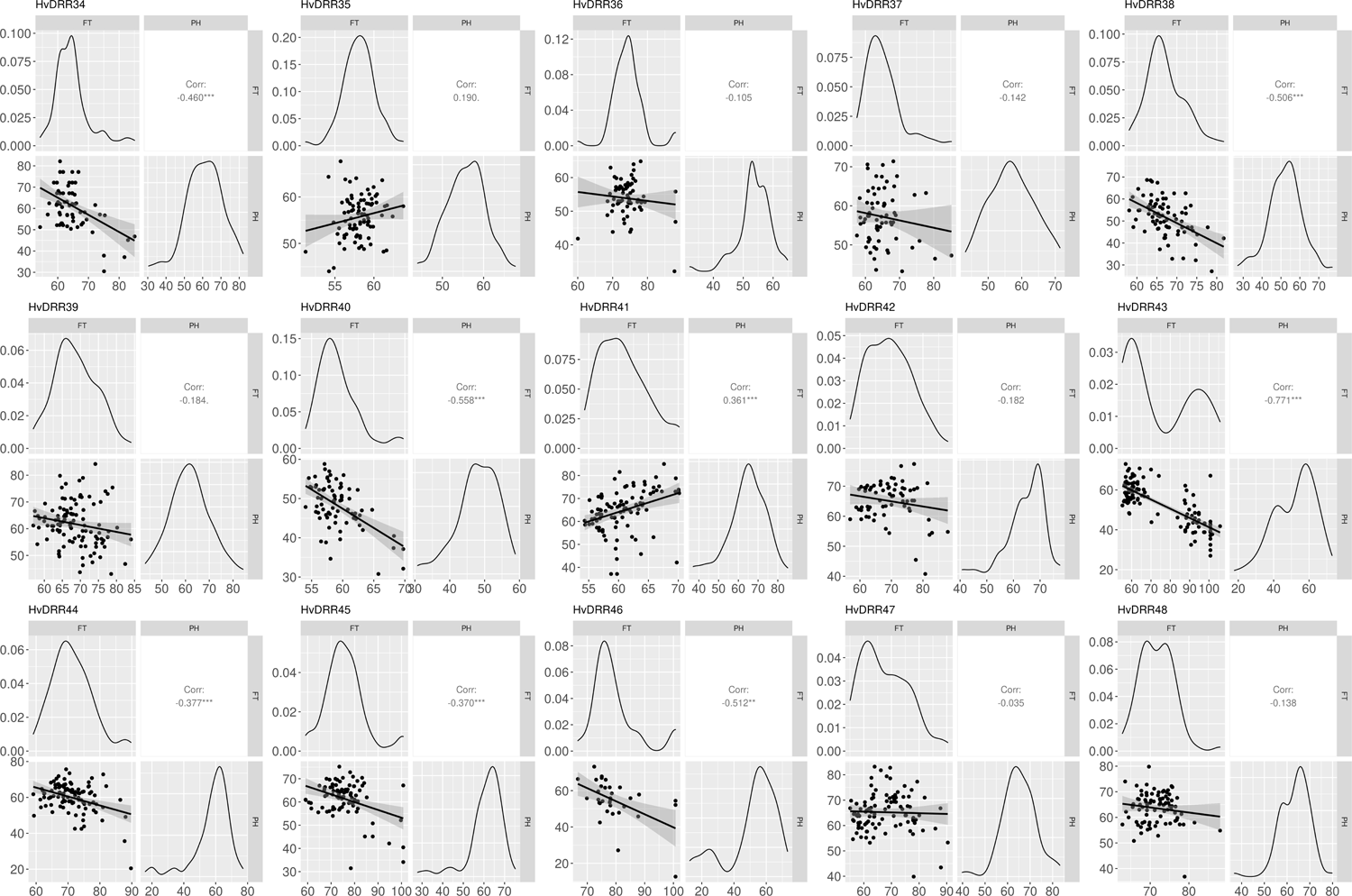
Histograms and correlation plots between flowering time (FT, in days after sowing) and plant height (PH, in cm), for each of the 45 HvDRR sub-populations.

**Supplementary Figure 3:**
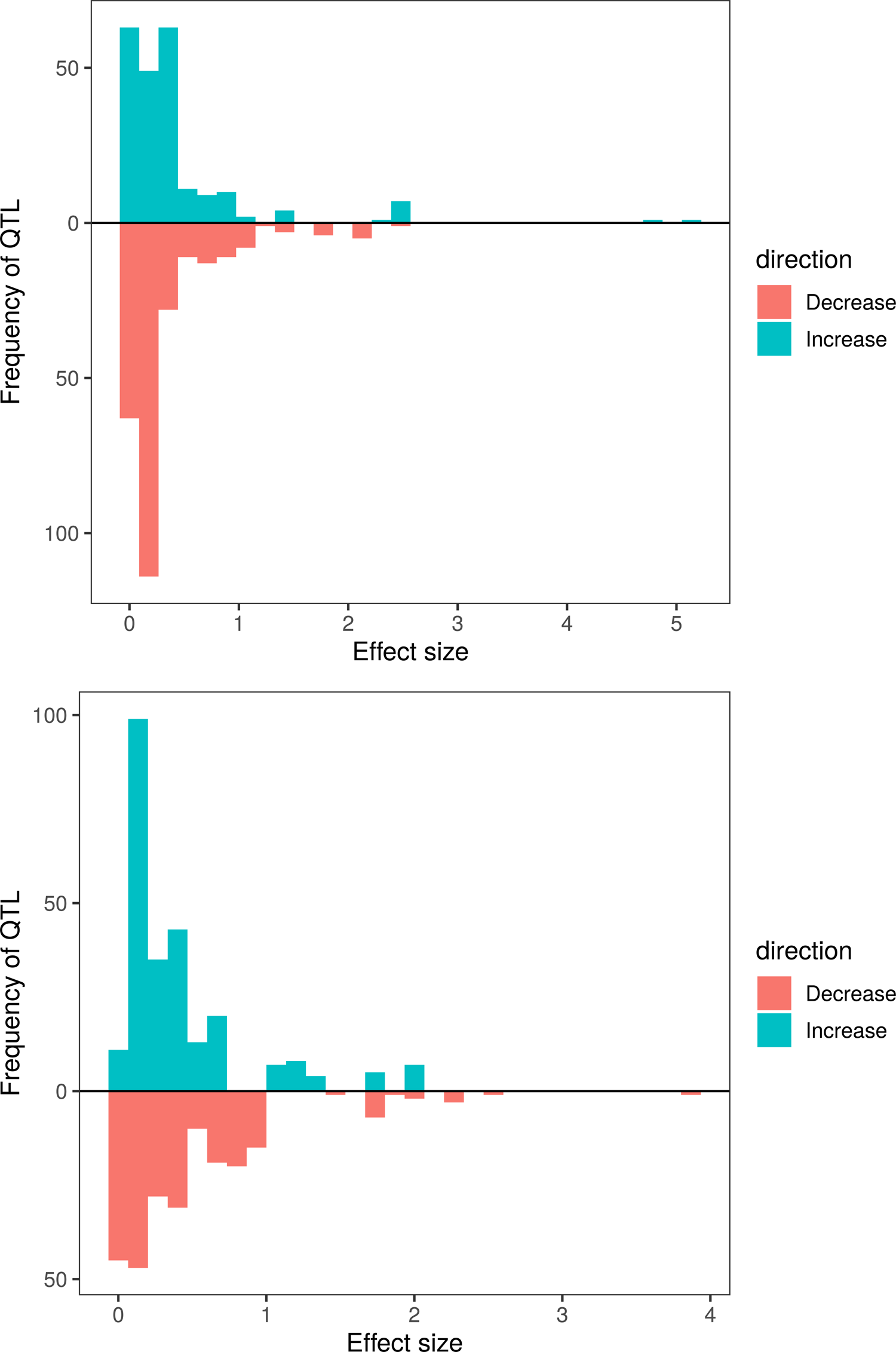
Effect size of the QTL detected through multi-parent population analysis for flowering time (top, in days after sowing) and plant height (bottom, in cm), for each of the parental lines.

**Supplementary Figure 4:**
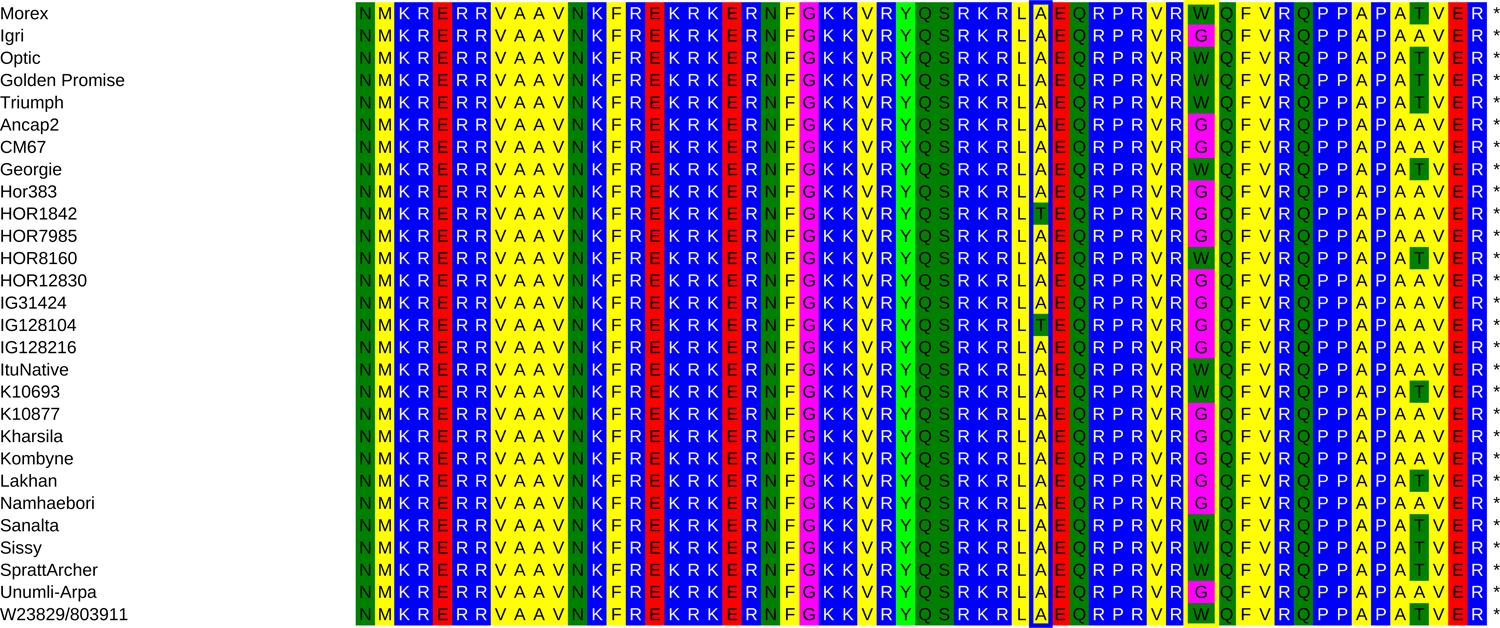
Amino acid sequence of the terminal region of *Ppd-H1* of Morex, Igri, Optic, Golden Promise, Triumph, and the 23 parental inbreds of the HvDRR population. The amino acid synthesized by the triplet containing SNP 22 is highlighted in yellow, the one synthesized by the triplet containing SNP 1945 is highlighted in blue.

**Supplementary Figure 5:**
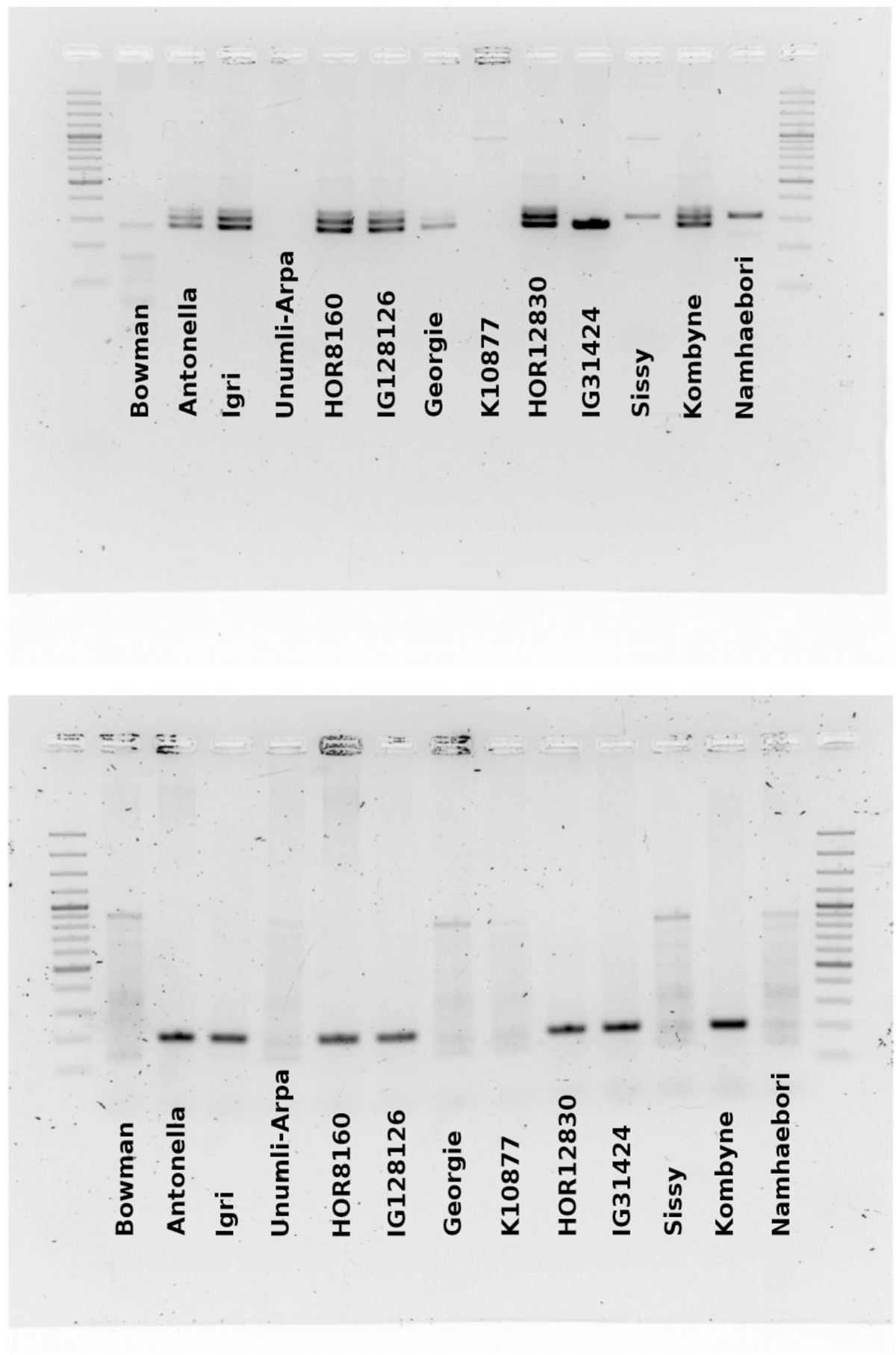
Gel pictures of PCRs performed to detect the presence/absence of *ZCCT-Ha:b* (top) and *ZCCT-Hc* (bottom) as described in Karsai *et al*. (2005). The analyzed genotypes are Bowman (control spring variety), Antonella (control winter variety), Igri (control winter variety), and the parental inbreds of the sub-populations for which a QTL co-localizing with *Vrn-H2* was detected.

**Supplementary Figure 6:**
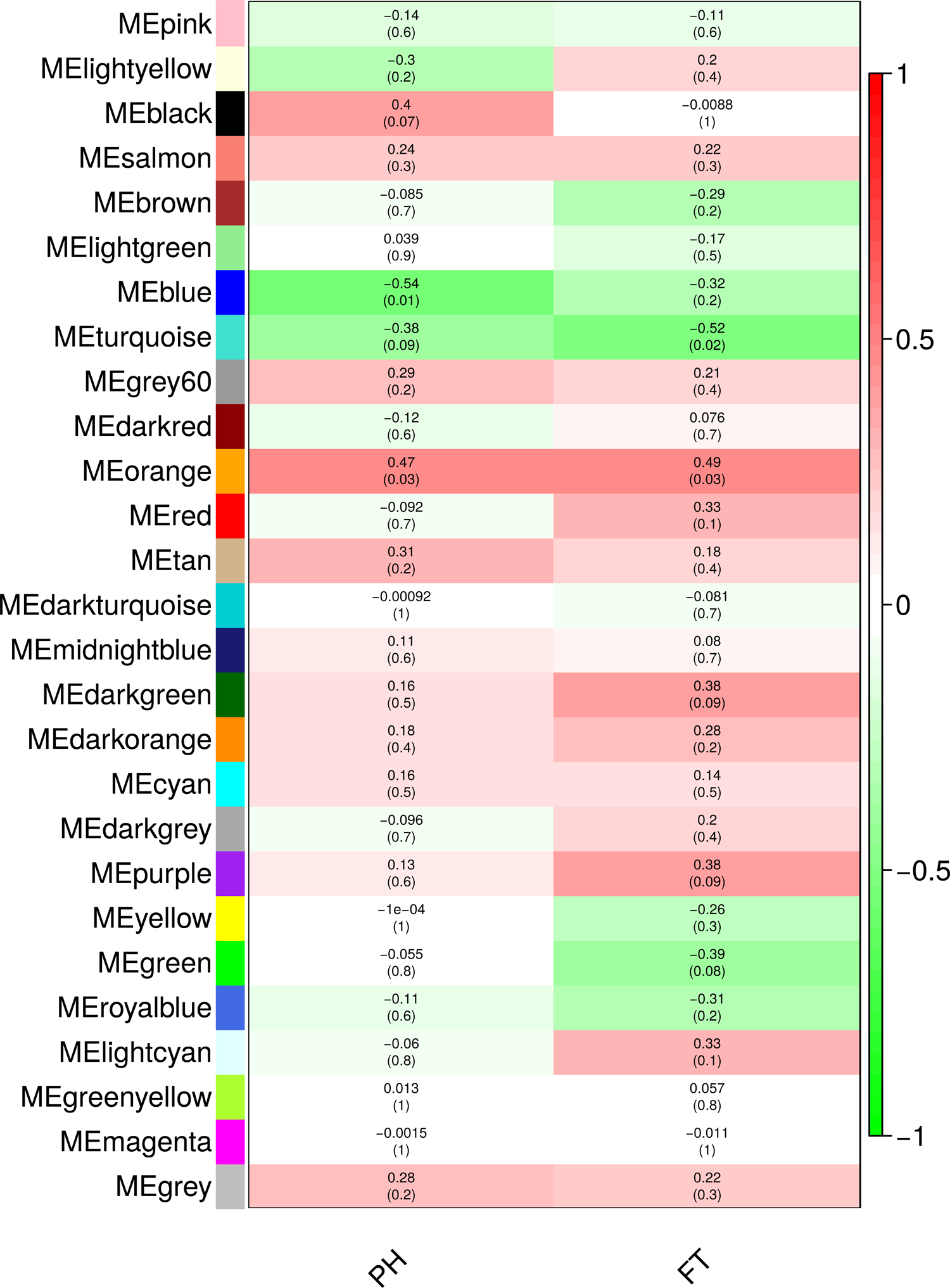
Heat map of the module-trait relationships for plant height (PH) and flowering time (FT). On the y axis, the 27 detected modules are reported. For each module-trait correlation p-values are given in brackets.

**Supplementary Figure 7:**
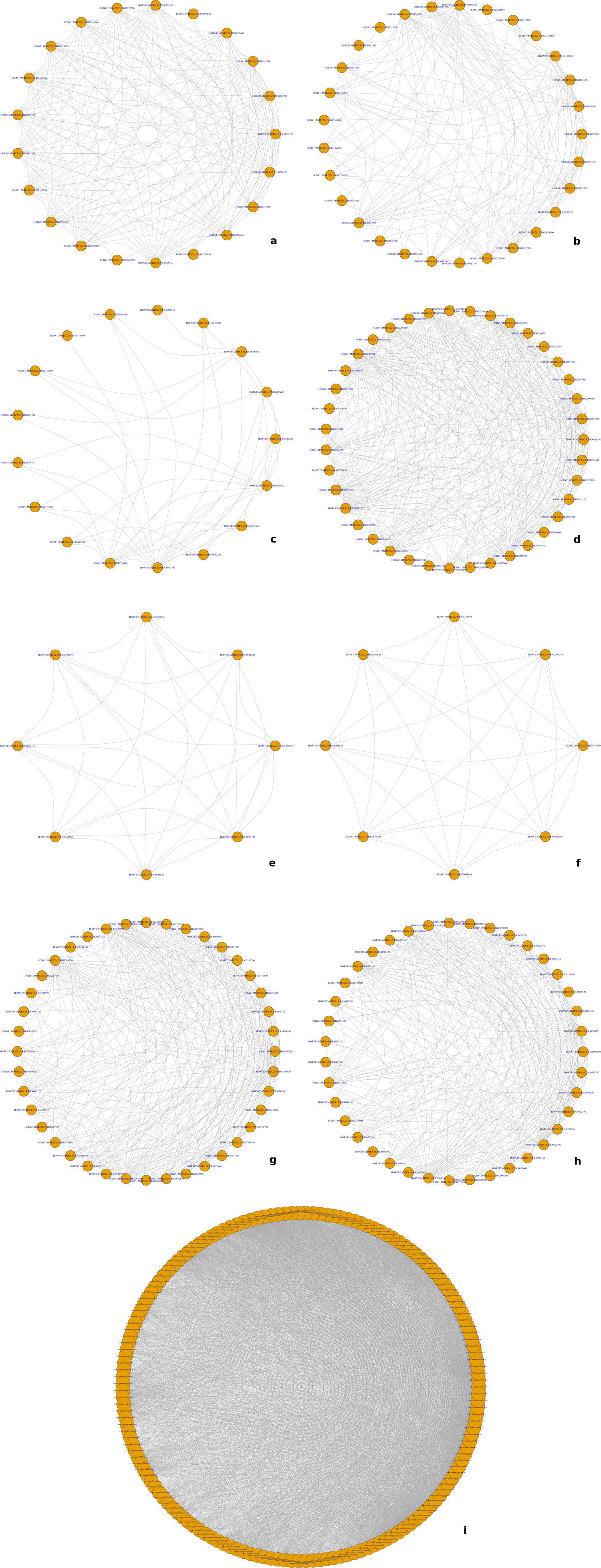
Network predictions for modules “orange” (a), “black” (b), “darkgreen” (c), “purple” (d), “tan” (e), “lightyellow” (f), “green” (g), “blue” (h), and “turquoise” (i). Gene names with a gene-module membership p-value < 0.01 are indicated in the orange circles. Gene-gene interactions are represented by grey lines.

**Supplementary Figure 8:**
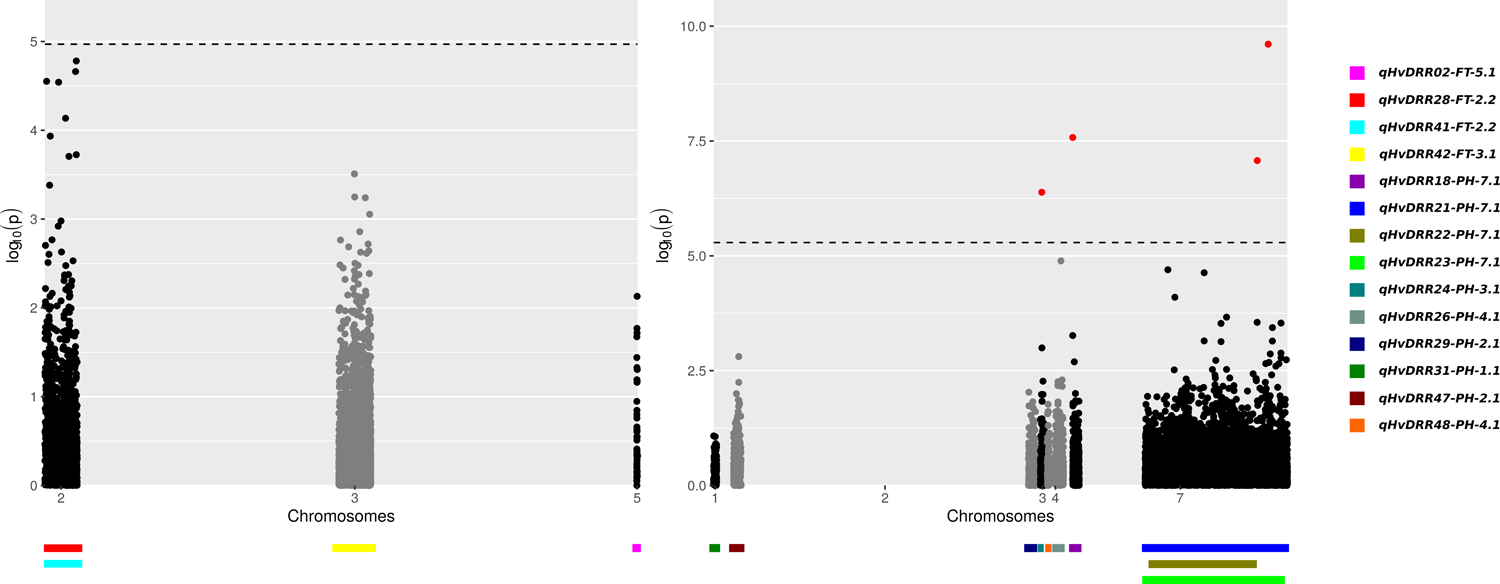
Negative decadic logarithm of the p-value for association tests of sequence variants in QTL without previously reported genes for the control of the trait within their interval, explaining ≥ 15% variance, and with interval ≤ 30 cM for flowering time (left) and plant height (right). The QTL confidence intervals from single population analyses are indicated by colored bars.

## REFERENCES

1. Afsharyan, N.P., Sannemann, W., Léon, J., and Ballvora, A. (2020) Effect of epistasis and environment on flowering time in barley reveals a novel flowering-delaying QTL allele. J. Exp. Bot., 71, 893–906.

2. Anderson, J.T. and Song, B.H. (2020) Plant adaptation to climate change—Where are we? J. Syst. Evol., 58, 533–545.

3. Araus, J.L., Slafer, G.A., Royo, C., and Serret, M.D. (2008) Breeding for yield potential and stress adaptation in cereals. CRC. Crit. Rev. Plant Sci., 27, 377–412.

4. Arifuzzaman, M., Günal, S., Bungartz, A., Muzammil, S., Afsharyan, N.P., Léon, J., and Naz, A.A. (2016) Genetic mapping reveals broader role of vrn-h3 gene in root and shoot development beyond heading in barley. PLoS One, 11, 1–16.

5. Arifuzzaman, M., Sayed, M.A., Muzammil, S., Pillen, K., Schumann, H., Naz, A.A., and Léon, J. (2014) Detection and validation of novel QTL for shoot and root traits in barley (Hordeum vulgare L.). Mol. Breed., 34, 1373–1387.

6. Bayer, M.M., Rapazote-Flores, P., Ganal, M., Hedley, P.E., Macaulay, M., Plieske, J., et al. (2017) Development and evaluation of a barley 50k iSelect SNP array. Front. Plant Sci., 8, 1–10.

7. Bezant, J., Laurie, D., Pratchett, N., Chojecki, J., and Kearsey, M. (1996) Marker regression mapping of QTL controlling flowering time and plant height in a spring barley (Hordeum vulgare L.) cross. Heredity (Edinb)., 77, 64–73.

8. Bi, X., Van Esse, W., Mulki, M.A., Kirschner, G., Zhong, J., Simon, R., and Von Korff, M. (2019) Centroradialis interacts with flowering locus t-like genes to control floret development and grain number. Plant Physiol., 180, 1013–1030.

9. Bleecker, A.B. and Kende, H. (2000) Ethylene: A gaseous signal molecule in plant. Annu. Rev. Cell Dev. Biol., 16, 1–18.

10. Broman, K.W., Wu, H., Sen, Ś., and Churchill, G.A. (2003) R/qtl: QTL mapping in experimental crosses. Bioinformatics, 19, 889–890.

11. Campoli, C., Drosse, B., Searle, I., Coupland, G., and Von Korff, M. (2012) Functional characterisation of HvCO1, the barley (Hordeum vulgare) flowering time ortholog of CONSTANS. Plant J., 69, 868–880.

12. Campoli, C., Pankin, A., Drosse, B., Casao, C.M., Davis, S.J., and Von Korff, M. (2013) HvLUX1 is a candidate gene underlying the early maturity 10 locus in barley: Phylogeny, diversity, and interactions with the circadian clock and photoperiodic pathways. New Phytol., 199, 1045–1059.

13. Casale, F., Van Inghelandt, D., Weisweiler, M., Li, J., and Stich, B. (2022) Genomic prediction of the recombination rate variation in barley – A route to highly recombinogenic genotypes. Plant Biotechnol. J., 20, 676–690.

14. Casao, M.C., Karsai, I., Igartua, E., Gracia, M.P., Veisz, O., and Casas, A.M. (2011) Adaptation of barley to mild winters: A role for PPDH2. BMC Plant Biol., 11.

15. Cockram, J., Jones, H., Leigh, F.J., O’Sullivan, D., Powell, W., Laurie, D.A., and Greenland, A.J. (2007) Control of flowering time in temperate cereals: Genes, domestication, and sustainable productivity. J. Exp. Bot., 58, 1231–1244.

16. Comadran, J., Kilian, B., Russell, J., Ramsay, L., Stein, N., Ganal, M., et al. (2012) Natural variation in a homolog of Antirrhinum CENTRORADIALIS contributed to spring growth habit and environmental adaptation in cultivated barley. Nat. Genet., 44, 1388–1391.

17. Cuesta-Marcos, A., Casas, A.M., Yahiaoui, S., Gracia, M.P., Lasa, J.M., and Igartua, E. (2008) Joint analysis for heading date QTL in small interconnected barley populations. Mol. Breed., 21, 383–399.

18. Dawson, I.K., Russell, J., Powell, W., Steffenson, B., Thomas, W.T.B., and Waugh, R. (2015) Barley: A translational model for adaptation to climate change. New Phytol., 206, 913–931.

19. Deng, W., Casao, M.C., Wang, P., Sato, K., Hayes, P.M., Finnegan, E.J., and Trevaskis, B. (2015) Direct links between the vernalization response and other key traits of cereal crops. Nat. Commun., 6.

20. Distelfeld, A., Li, C., and Dubcovsky, J. (2009) Regulation of flowering in temperate cereals. Curr. Opin. Plant Biol., 12, 178–184.

21. Dockter, C., Gruszka, D., Braumann, I., Druka, A., Druka, I., Franckowiak, J., et al. (2014) Induced variations in brassinosteroid genes define barley height and sturdiness, and expand the green revolution genetic toolkit. Plant Physiol., 166, 1912–1927.

22. Druka, A., Franckowiak, J., Lundqvist, U., Bonar, N., Alexander, J., Houston, K., et al. (2011) Genetic dissection of barley morphology and development. Plant Physiol., 155, 617–627.

23. Dubois, M., Van den Broeck, L., and Inzé, D. (2018) The Pivotal Role of Ethylene in Plant Growth. Trends Plant Sci., 23, 311–323.

24. Dunford, R.P., Griffiths, S., Christodoulou, V., and Laurie, D.A. (2005) Characterisation of a barley (Hordeum vulgare L.) homologue of the Arabidopsis flowering time regulator GIGANTEA. Theor. Appl. Genet., 110, 925–931.

25. Eriksson, M.E. and Millar, A.J. (2003) The circadian clock. A plant’s best friend in a spinning world. Plant Physiol., 132, 732–738.

26. Food and Agriculture Organization of the United Nations. (2019). FAOSTAT statistical database. [Rome]: FAO

27. Food and Agriculture Organization of the United Nations. (2020). FAOSTAT statistical database. [Rome]: FAO

28. Faure, S., Turner, A.S., Gruszka, D., Christodoulou, V., Davis, S.J., Von Korff, M., and Laurie, D.A. (2012) Mutation at the circadian clock gene EARLY MATURITY 8 adapts domesticated barley (Hordeum vulgare) to short growing seasons. Proc. Natl. Acad. Sci. U. S. A., 109, 8328–8333.

29. Fernández-Calleja, M., Casas, A.M., and Igartua, E. (2021) Major flowering time genes of barley: allelic diversity, effects, and comparison with wheat. Springer Berlin Heidelberg.

30. Garin, V., Wimmer, V., and Malosetti, M. (2015) mppR : An R Package for QTL Analysis in Multi-parent Populations using Linear Mixed Models.

31. Garin, V., Wimmer, V., Mezmouk, S., Malosetti, M., and van Eeuwijk, F. (2017) How do the type of QTL effect and the form of the residual term influence QTL detection in multi-parent populations? A case study in the maize EU-NAM population. Theor. Appl. Genet., 130, 1753–1764.

32. Giraud, H., Lehermeier, C., Bauer, E., Falque, M., Segura, V., Bauland, C., et al. (2014) Linkage disequilibrium with linkage analysis of multiline crosses reveals different multiallelic QTL for hybrid performance in the flint and dent heterotic groups of maize. Genetics, 198, 1717–1734.

33. Göransson, M., Hallsson, J.H., Lillemo, M., Orabi, J., Backes, G., Jahoor, A., et al. (2019) Identification of ideal allele combinations for the adaptation of spring barley to northern latitudes. Front. Plant Sci., 10, 1–13.

34. Hada, W., Bo, Z., Cao, W.H., Biao, M., Gang, L., Liu, Y.F., et al. (2009) The Ethylene Receptor ETR2 delays floral transition and affects starch accumulation in rice. Plant Cell, 21, 1473–1494.

35. Hayama, R. and Coupland, G. (2004) The molecular basis of diversity in the photoperiodic flowering responses of arabidopsis and rice. Plant Physiol., 135, 677–684.

36. Hemming, M.N., Peacock, W.J., Dennis, E.S., and Trevaskis, B. (2008) Low-temperature and daylength cues are integrated to regulate Flowering Locus T in barley. Plant Physiol., 147, 355–366.

37. Hemshrot, A., Poets, A.M., Tyagi, P., Lei, L., Carter, C.K., Hirsch, C.N., et al. (2019) Development of a multiparent population for genetic mapping and allele discovery in six-row barley. Genetics, 213, 595–613.

38. Hill, C.B. and Li, C. (2016) Genetic architecture of flowering phenology in cereals and opportunities for crop improvement. Front. Plant Sci., 7, 1–23.

39. Iqbal, N., Khan, N.A., Ferrante, A., Trivellini, A., Francini, A., and Khan, M.I.R. (2017) Ethylene role in plant growth, development and senescence: interaction with other phytohormones. Front. Plant Sci., 8, 1–19.

40. Jia, Q.J., Zhang, J.J., Westcott, S., Zhang, X.Q., Bellgard, M., Lance, R., and Li, C.D. (2009) GA-20 oxidase as a candidate for the semidwarf gene sdw1/denso in barley. Funct. Integr. Genomics, 9, 255–262.

41. Jones, H., Leigh, F.J., Mackay, I., Bower, M.A., Smith, L.M.J., Charles, M.P., et al. (2008) Population-based resequencing reveals that the flowering time adaptation of cultivated barley originated east of the fertile crescent. Mol. Biol. Evol., 25, 2211– 2219.

42. Kang, H.M., Sul, J.H., Service, S.K., Zaitlen, N.A., Kong, S.Y., Freimer, N.B., et al. (2010) Variance component model to account for sample structure in genome-wide association studies. Nat. Genet., 42, 348–354.

43. Karsai, I., Szucs, P., Mészáros, K., Filichkina, T., Hayes, P.M., Skinner, J.S., et al. (2005) The Vrn-H2 locus is a major determinant of flowering time in a facultative x winter growth habit barley (Hordeum vulgare L.) mapping population. Theor. Appl. Genet., 110, 1458–1466.

44. Khush, G.S. (2013) Strategies for increasing the yield potential of cereals: Case of rice as an example. Plant Breed., 132, 433–436.

45. Kikuchi, R. and Handa, H. (2009) Photoperiodic control of flowering in barley. Breed. Sci., 59, 546–552.

46. Knott, S.A. and Haley, C.S. (1992) A simple regression method for mapping quantitative trait loci in line crosses using flanking markers. Heredity (Edinb)., 69, 315–324.

47. Von Korff, M., Wang, H., Léon, J., and Pillen, K. (2006) AB-QTL analysis in spring barley: II. Detection of favourable exotic alleles for agronomic traits introgressed from wild barley (H. vulgare ssp. spontaneum). Theor. Appl. Genet., 112, 1221–1231.

48. Langfelder, P. and Horvath, S. (2008) WGCNA: An R package for weighted correlation network analysis. BMC Bioinformatics, 9.

49. Langridge (2018), Peter. Economic and academic importance of barley. In The barley genome. Springer, Cham, 1-10.

50. Laurie, D.A., Pratchett, N., Bezant, J.H., and Snape, J.W. (1994) Genetic analysis of a photoperiod response gene on the short arm of chromosome 2(2h) of Hordeum vulgare (barley). Heredity (Edinb)., 72, 619–627.

51. Li, Z.K., Yu, S.B., Lafitte, H.R., Huang, N., Courtois, B., Hittalmani, S., et al. (2003) QTL x environment interactions in rice. I. Heading date and plant height. Theor. Appl. Genet., 108, 141–153.

52. Manichaikul, A., Dupuis, J., Sen, Ś., and Broman, K.W. (2006) Poor performance of bootstrap confidence intervals for the location of a quantitative trait locus. Genetics, 174, 481–489.

53. Mascher, M., Wicker, T., Jenkins, J., Plott, C., Lux, T., Koh, C.S., et al. (2021) Long-read sequence assembly: A technical evaluation in barley. Plant Cell, 33, 1888–1906.

54. Matsushika, A., Makino, S., Kojima, M., and Mizuno, T. (2000) Circadian waves of expression of the APRR1/TOC1 family of pseudo-response regulators in Arabidopsis thaliana: Insight into the plant circadian clock. Plant Cell Physiol., 41, 1002–1012.

55. Maurer, A., Draba, V., Jiang, Y., Schnaithmann, F., Sharma, R., Schumann, E., et al. (2015) Modelling the genetic architecture of flowering time control in barley through nested association mapping. BMC Genomics, 16, 1–12.

56. Maurer, A., Draba, V., and Pillen, K. (2016) Genomic dissection of plant development and its impact on thousand grain weight in barley through nested association mapping. J. Exp. Bot., 67, 2507–2518.

57. Mikołajczak, K., Kuczyńska, A., Krajewski, P., Sawikowska, A., Surma, M., Ogrodowicz, P., et al. (2017) Quantitative trait loci for plant height in Maresi × CamB barley population and their associations with yield-related traits under different water regimes. J. Appl. Genet., 58, 23–35.

58. Sharma, R., Shaaf, S., Neumann, K., Go, Y., Mascher, M., David, M., et al. (2020) On the origin of photoperiod non-responsiveness in barley. bioRxiv, 2020.07.02.185488.

59. Mizuno, T. and Nakamichi, N. (2005) Pseudo-response regulators (PRRs) or true oscillator components (TOCs). Plant Cell Physiol., 46, 677–685.

60. Mulki, M.A. and von Korff, M. (2016) CONSTANS controls floral repression by up-regulating VERNALIZATION2 (VRN-H2) in Barley1. Plant Physiol., 170, 325–337.

61. Myles, S., Peiffer, J., Brown, P.J., Ersoz, E.S., Zhang, Z., Costich, D.E., and Buckler, E. (2009) Association mapping: Critical considerations shift from genotyping to experimental design. Plant Cell, 21, 2194–2202.

62. Nakamichi, N., Kudo, T., Makita, N., Kiba, T., Kinoshita, T., and Sakakibara, H. (2020) Flowering time control in rice by introducing Arabidopsis clock-associated PSEUDO-RESPONSE REGULATOR 5. Biosci. Biotechnol. Biochem., 84, 970–979.

63. Nice, L.M., Steffenson, B.J., Blake, T.K., Horsley, R.D., Smith, K.P., and Muehlbauer, G.J. (2017) Mapping agronomic traits in a wild barley advanced backcross–nested association mapping population. Crop Sci., 57, 1199–1210.

64. Nishida, H., Ishihara, D., Ishii, M., Kaneko, T., Kawahigashi, H., Akashi, Y., et al. (2013) Phytochrome C is a key factor controlling long-day flowering in barley. Plant Physiol., 163, 804–814.

65. Pasam, R.K., Sharma, R., Malosetti, M., van Eeuwijk, F. a, Haseneyer, G., Kilian, B., and Graner, A. (2012) Genome-wide association studies for agronomical traits in a world wide spring barley collection. BMC Plant Biol., 12, 16.

66. Patil, V., McDermott, H.I., McAllister, T., Cummins, M., Silva, J.C., Mollison, E., et al. (2019) APETALA2 control of barley internode elongation. Dev., 146.

67. Pauli, D., Muehlbauer, G.J., Smith, K.P., Cooper, B., Hole, D., Obert, D.E., et al. (2014) Association Mapping of Agronomic QTLs in U.S. Spring Barley Breeding Germplasm. Plant Genome, 7.

68. Pieper, R., Tomé, F., Pankin, A., and Von Korff, M. (2021) FLOWERING LOCUS T4 delays flowering and decreases floret fertility in barley. J. Exp. Bot., 72, 107–121.

69. Piepho, H.P. and Möhring, J. (2007) Computing heritability and selection response from unbalanced plant breeding trials. Genetics, 177, 1881–1888.

70. Rodriguez, M., Rau, D., Papa, R., and Attene, G. (2008) Genotype by environment interactions in barley (Hordeum vulgare L.): Different responses of landraces, recombinant inbred lines and varieties to Mediterranean environment. Euphytica, 163, 231–247.

71. Rollins, J.A., Drosse, B., Mulki, M.A., Grando, S., Baum, M., Singh, M., et al. (2013) Variation at the vernalisation genes Vrn-H1 and Vrn-H2 determines growth and yield stability in barley (Hordeum vulgare) grown under dryland conditions in Syria. Theor. Appl. Genet., 126, 2803–2824.

72. van Rossum, B.J., Kruijer, W., van Eeuwijk, F., Boer, M., Malosetti, M., Bustos-Korts, D., and Wehrens, R. (2022) Package ‘ statgenGWAS *.’* R Packag. version, 1.

73. Schmalenbach, I., Léon, J., and Pillen, K. (2009) Identification and verification of QTLs for agronomic traits using wild barley introgression lines. Theor. Appl. Genet., 118, 483– 497.

74. Sharma, R., Shaaf, S., Neumann, K., Go, Y., Mascher, M., David, M., et al. (2020) On the origin of photoperiod non-responsiveness in barley. bioRxiv, 2020.07.02.185488.

75. Shoesmith, J.R., Solomon, C.U., Yang5, X., Wilkinson, L.G., Sheldrick, S., Van Eijden, E., et al. (2021) APETALA2 functions as a temporal factor together with BLADE-ON-PETIOLE2 and MADS29 to control flower and grain development in barley. Dev., 148.

76. Shrestha, A., Cosenza, F., van Inghelandt, D., Wu, P.-Y., Li, J., Casale, F.A., et al. (2022) The double round-robin population unravels the genetic architecture of grain size in barley. J. Exp. Bot., 73, 7344–7361.

77. Stich, B. (2009) Comparison of mating designs for establishing nested association mapping populations in maize and Arabidopsis thaliana. Genetics, 183, 1525–1534.

78. Turner, A., Beales, J., Faure, S., Dunford, R.P., and Laurie, D.A. (2005) The pseudo-response regulator Ppd-H1 provides adaptation to photoperiod in barley. Science (80-.)., 310, 1031–1034.

79. VanRaden, P.M. (2008) Efficient methods to compute genomic predictions. J. Dairy Sci., 91, 4414–4423.

80. Vaser, R., Adusumalli, S., Leng, S.N., Sikic, M., and Ng, P.C. (2016) SIFT missense predictions for genomes. Nat. Protoc., 11, 1–9.

81. Vidal, T., Gigot, C., De Vallavieille-Pope, C., Huber, L., and Saint-Jean, S. (2018) Contrasting plant height can improve the control of rain-borne diseases in wheat cultivar mixture: Modelling splash dispersal in 3-D canopies. Ann. Bot., 121, 1299– 1308.

82. Vyas, S., Khatri-Chhetri, A., Aggarwal, P., Thornton, P., and Campbell, B.M. (2022) Perspective: The gap between intent and climate action in agriculture. Glob. Food Sec., 32, 100612.

83. Wang, J., Yang, J., Jia, Q., Zhu, J., Shang, Y., Hua, W., and Zhou, M. (2014) A new QTL for plant height in barley (hordeum vulgare l.) showing no negative effects on grain yield. PLoS One, 9.

84. Wei, J., Fang, Y., Jiang, H., ting Wu, X., hong Zuo, J., chun Xia, X., et al. (2022) Combining QTL mapping and gene co-expression network analysis for prediction of candidate genes and molecular network related to yield in wheat. BMC Plant Biol., 22, 1–14.

85. Weisweiler, M., Arlt, C., Wu, P.-Y., Van Inghelandt, D., Hartwig, T., Stich, B., et al. (2022) Structural variants in the barley gene pool: precision and sensitivity to detect them using short-read sequencing and their association with gene expression and phenotypic variation. bioRxiv, 2022.04.25.489331.

86. Weisweiler, M., De Montaigu, A., Ries, D., Pfeifer, M., and Stich, B. (2019) Transcriptomic and presence/absence variation in the barley genome assessed from multi-tissue mRNA sequencing and their power to predict phenotypic traits. BMC Genomics, 20, 1–15.

87. Wendt, T., Holme, I., Dockter, C., Preu, A., Thomas, W., Druka, A., et al. (2016) HvDep1 Is a Positive regulator of culm elongation and grain size in barley and impacts yield in an environment-dependent manner. PLoS One, 11, 1–21.

88. Wiegmann, M., Maurer, A., Pham, A., March, T.J., Al-Abdallat, A., Thomas, W.T.B., et al. (2019) Barley yield formation under abiotic stress depends on the interplay between flowering time genes and environmental cues. Sci. Rep., 9, 1–16.

89. Yan, L., Fu, D., Li, C., Blechl, A., Tranquilli, G., Bonafede, M., et al. (2006) The wheat and barley vernalization gene VRN3 is an orthologue of FT. Proc. Natl. Acad. Sci. U. S. A., 103, 19581–19586.

90. Yan, L., Loukoianov, A., Blechl, A., Tranquilli, G., Ramakrishna, W., SanMiguel, P., et al. (2004) The Wheat VRN2 Gene Is a Flowering Repressor Down-Regulated by Vernalization. Science (80-.)., 303, 1640–1644.

91. Yan, L., Loukoianov, A., Tranquilli, G., Helguera, M., Fahima, T., and Dubcovsky, J. (2003) Positional cloning of the wheat vernalization gene VRN1. Proc. Natl. Acad. Sci. U. S. A., 100, 6263–6268.

92. Yu, J., Pressoir, G., Briggs, W.H., Bi, I.V., Yamasaki, M., Doebley, J.F., et al. (2006) A unified mixed-model method for association mapping that accounts for multiple levels of relatedness. Nat. Genet., 38, 203–208.

93. Zakhrabekova, S., Gough, S.P., Braumann, I., Muller, A.H., Lundqvist, J., Ahmann, K., et al. (2012) Induced mutations in circadian clock regulator Mat-a facilitated short-season adaptation and range extension in cultivated barley. Proc. Natl. Acad. Sci. U. S. A., 109, 4326–4331.

94. Zhang, B. and Horvath, S. (2005) A general framework for weighted gene co-expression network analysis. Stat. Appl. Genet. Mol. Biol., 4.

